# Defective pgsA contributes to increased membrane fluidity and cell wall thickening in S. aureus with high-level daptomycin resistance

**DOI:** 10.1101/2023.04.11.536441

**Authors:** Christian D. Freeman, Tayte Hansen, Ramona Urbauer, Brian J. Wilkinson, Vineet K. Singh, Kelly M. Hines

## Abstract

Daptomycin is a membrane-targeting last-resort antimicrobial therapeutic for the treatment of infections caused by methicillin- and/or vancomycin-resistant *Staphylococcus aureus*. In the rare event of failed daptomycin therapy, the source of resistance is often attributable to mutations directly within the membrane phospholipid biosynthetic pathway of *S. aureus* or in the regulatory systems that control cell envelope response and membrane homeostasis. Here we describe the structural changes to the cell envelope in a daptomycin-resistant isolate of *S. aureus* strain N315 that has acquired mutations in the genes most commonly reported associated with daptomycin-resistance: *mprF*, *yycG*, and *pgsA*. In addition to the decreased phosphatidylglycerol (PG) levels that are the hallmark of daptomycin-resistance, the mutant with high-level daptomycin resistance had increased branched-chain fatty acids (BCFAs) in its membrane lipids, increased membrane fluidity, and increased cell wall thickness. However, the successful utilization of isotope-labeled straight-chain fatty acids (SCFAs) in lipid synthesis suggested that the aberrant BCFA:SCFA ratio arose from upstream alteration in fatty acid synthesis rather than a structural preference in PgsA. RT-qPCR studies revealed that expression of pyruvate dehydrogenase (*pdhB*) was suppressed in the daptomycin-resistant isolate, which is known to increase BCFA levels. While complementation with an additional copy of *pdhB* had no effect, complementation of the *pgsA* mutation resulted in increased PG formation, reduction in cell wall thickness, restoration of normal BCFA levels, and increased daptomycin susceptibility. Collectively, these results demonstrate that *pgsA* contributes to daptomycin resistance through its influence on membrane fluidity and cell wall thickness, in addition to phosphatidylglycerol levels.

**IMPORTANCE:** The cationic lipopeptide antimicrobial daptomycin has become an essential tool for combating infections with *Staphylococcus aureus* that display reduced susceptibility to β-lactams or vancomycin. Since daptomycin’s activity is based on interaction with the negatively charged membrane of *S. aureus*, routes to daptomycin-resistance occur through mutations in the lipid biosynthetic pathway surrounding phosphatidylglycerols and the regulatory systems that control cell envelope homeostasis. Therefore, there are many avenues to achieve daptomycin resistance and several different, and sometimes contradictory, phenotypes of daptomycin-resistant *S. aureus,* including both increased and decreased cell wall thickness and membrane fluidity. This study is significant because it demonstrates the unexpected influence of a lipid biosynthesis gene, *pgsA*, on membrane fluidity and cell wall thickness in *S. aureus* with high-level daptomycin resistance.

## INTRODUCTION

Whereas many of the first-choice antimicrobials, including β-lactams and vancomycin, for the treatment of infections with Gram-positive bacteria inhibit cell wall biosynthesis, the cationic lipopeptide antibiotic daptomycin targets the negatively-charged membrane of bacteria and, through mechanisms that are still debated, causes cell death by altering the integrity and structure of the cell envelope (1–6). The differences between the mechanisms of daptomycin and β-lactams/vancomycin have led to its use as a “last-resort” therapeutic in situations when resistance has established against one or more cell wall-targeting antimicrobials (7, 8). The growing incidences of methicillin-resistant *S. aureus* (MRSA) and vancomycin intermediate *S. aureus* (VISA) will likely increase the utilization of daptomycin, providing more opportunities for daptomycin resistance to emerge.

Common phenotypes of daptomycin-resistant MRSA include thickened cell walls, a reduction in the negative cell surface charge, and altered membrane fluidity (9–13). These changes to the organism’s cell envelope are the result of mutations in regulatory and synthetic pathways for cell wall synthesis, membrane homeostasis, and lipid biosynthesis (13–16). The cell wall thickening is attributed to mutations in the *vraSR* or *yycFG* (or *walKR*) two-component regulatory systems that respond to stimuli by promoting transcription of genes encoding for proteins that synthesize, metabolize, or otherwise modify components of the cell wall (15–17). Increased modification of wall teichoic acids with D-alanine due to upregulation of the *dlt* operon has also been reported in Dap-R isolates (18–20). However, thickened cell walls and increased D-alanylation are not universal phenotypes for Dap-R in *S. aureus* (21).

The key mechanisms that promote daptomycin resistance occur within the membrane and specifically affect the amount of the negatively-charged phosphatidylglycerols (PG) that are targeted by the cationic daptomycin molecules. The most common genetic route to decreasing PGs detected in clinical isolates is mutation to the gene encoding MprF, a lysyl-phosphatidylglycerol (LysylPG) synthase and flippase (22–24). The addition of a lysine moiety to the headgroup of PG effectively converts the negatively-charged phospholipid into a cationic species. Translocation of the LysylPGs to the outer membrane leaflet decreases the net-negative charge of the membrane surface and reduces the affinity of daptomycin binding (24). Other mechanisms to reduce PG levels include mutations in phosphatidylglycerol synthase, *pgsA*, which reduces PG synthesis, or mutations in cardiolipin synthase, *cls2*, that promote the turnover of PGs into cardiolipins (CLs) (25–27). However, daptomycin-resistant isolates of *S. aureus* with *cls2* and *pgsA* mutations are not commonly detected in clinical settings (26).

A previous study of daptomycin-resistance in *S. aureus* strain N315 found that high-level resistance (MIC = 8 µg/mL) was achieved by an *in vitro* serial passage mutant through a combination of the above-mentioned mutations: *mprF*, *yycG*, and *pgsA* (28). Lipidomics analyses performed by hydrophilic interaction liquid chromatography coupled to mass spectrometry (LC-MS) identified reductions in total levels of PGs, LysylPGs, and CLs. However, the patterns of individual lipid species revealed heterogeneity in the net reduction that was dependent on the acyl tail composition. This trend was more pronounced in lipids upstream in the biosynthetic pathway from PGs, including phosphatidic acids (PAs) and diglycosyldiacylglycerols (DGDGs), that were elevated due to the bottleneck of the *pgsA* mutation. Analysis of the fatty acid content of PGs and DGDGs, combined with the levels of free fatty acids (FFAs), indicated that odd-carbon and long-chain fatty acids (>18 carbons) were elevated in the daptomycin-resistant mutant. Therefore, it was suggested by Hines et al. that strain N315-D8 may have aberrantly high levels of branched-chain fatty acids and increased membrane fluidity (28).

We recently demonstrated a high-resolution chromatography method based on reversed-phase chromatography (RPLC) that can separate diacyl lipids (*e.g.*, PGs, LysylPGs, etc.) based on the number of branched vs. straight-chain fatty acids that they contain (29). Using this method, we set out to re-evaluate the lipidome of *S. aureus* strain N315-D8 with high-level daptomycin resistance for indications of altered BCFA production. Our results revealed that N315-D8 did in fact contain a substantially greater amount of BCFAs and that their presence promoted a highly fluid membrane in the organism. The lipidomics data also identified clear differences in the utilization of BCFAs vs. SCFAs between phospholipid species. Additional phenotypic studies determined that a thickened cell wall and increased negative surface charge also contribute to the daptomycin resistance of N315-D8. However, we have demonstrated that some of these phenotypes can be reversed in the Dap-R strain through complementation of the *pgsA* mutation and, to a lesser extent, incorporation of exogenous straight-chain fatty acids. The results of this study suggest that traits of daptomycin susceptible *S. aureus* can be restored in resistant strains by modulating the fatty acyl content of the membrane.

## METHODS

### Bacteria Strains and Growth Conditions

All work with *S. aureus* was performed with Biosafety Level 2 (BSL-2) precautions. Details on the strains used in this study are summarized in **Table 1**. *S. aureus* strains N315 (MRSA) and N315-D8 were streaked from frozen stocks directly onto sterile brain heart infusion (BHI) agar and incubated at 37°C overnight (28). Individual colonies were used to prepare 2 McFarland suspensions (∼6 x 10^8^ CFU/mL) in sterile saline (DEN-1B densitometer). Liquid cultures were prepared in 15 mL centrifuge tubes by inoculating 4.5 mL of sterile tryptic soy broth (TSB, BD Bacto) with 0.5 mL of the 2 McFarland suspensions. For fatty acid labeling experiments, 10 µL of 50 mM methyl-*d*_3_ pentadecanoic acid (98%, Cambridge Isotope Laboratories) or 7,7,8,8-*d_4_* palmitic acid (98%, Cambridge Isotope Laboratories) in ethanol was added into the liquid culture to a final concentration of 100 µM. Cultures containing 10 µL of ethanol were prepared as a vehicle control and a second control was prepared with supplemented broth. All conditions were prepared in triplicate. Liquid cultures were incubated at 37 °C with shaking (200 rpm) for 24 hrs, unless otherwise noted, to allow both N315 and N315-D8 to reach stationary phase (see **SI Figure S1**). Bacteria were harvested by centrifugation (3,000 rpm for 10 min at 4°C) and the resulting pellets were washed once with sterile water.

### Lipid Extraction

A modified version of the Bligh and Dyer method was used for the lipid extraction (30). Bacteria pellets were resuspended in LC-MS grade water (Optima, Fisher) to achieve McFarland readings differing by 0.04 units or less. A 2 mL portion of each suspension was transferred to glass centrifuge tubes. Samples were sonicated in an ice bath for 30 min. A chilled solution of 2:1 methanol/chloroform (v/v, 2mL) was added to each tube. After 5 min of periodic vortexing, 0.5 mL each of chilled chloroform and water were added to the tubes. Each sample was vortexed for one minute and centrifuged at 4 °C and 2,000 rpm for 10 min to induce phase separation. The organic layer containing the lipids was collected into fresh glass tubes, dried in a speed-vac, reconstituted with 0.5 mL of 1:1 chloroform and methanol (v/v). Samples were stored at −80 °C in sealed glass vials until analysis. For lipidomics experiments, 40 µL aliquots of each lipid extract were transferred to new vials, dried in a speed-vac, and reconstituted with 100 µL of MPA for negative mode (2.5x dilution) or 40 µL for positive mode (1x dilution).

### Lipidomics Analysis

Lipid separations were performed on a Waters Acquity FTN I-Class Plus ultra-performance liquid chromatography (UPLC) system equipped with a Waters Acquity charged surface hybrid (CSH) C18 (2.1 x 100mm, 1.7µm) column, as described previously (29). Mobile phase A (MPA) consisted of 60:40 acetonitrile/water with 10 mM ammonium formate. Mobile phase B (MPB) consisted of 88:10:2 isopropanol/acetonitrile/water with 10 mM ammonium formate. LysylPGs were analyzed separately using mobile phase solutions adjusted to pH 9.7 with ammonium hydroxide. All solvents and salts were LC-MS grade (Fisher Optima). A 30 min gradient elution was performed with a flow rate of 0.3 mL/min using the following conditions: 0-3 min, 30% MPB; 3-5 min, 30-50% MPB; 5-15 min, 50-90% MPB; 15-16 min, 90-99% MPB; 16-20 min, 99% MPB; 20-22 min, 99-30% MPB; 22-30 min 30% MPB. The column temperature was maintained at 40 °C and samples were kept at 6 °C. An injection volume of 10 µL was used.

The Waters Acquity UPLC was connected to the electrospray ionization source of a Waters Synapt XS traveling-wave ion mobility-mass spectrometer (TWIM-MS). Negative and positive mode ionizations were conducted using the following: Capillary voltage: +/-2.0 kV; sampling cone voltage: 40 V; source offset: 4V; source temperature: 120°C; desolvation temperature: 450°C; desolvation gas flow: 700 L/Hr; cone gas flow: 50 L/Hr. Traveling wave separations were done with a wave velocity of 550 m/s, wave height of 40 V, and a gas flow of 90mL/min (nitrogen). The time-of-flight mass analyzer operated in V-mode (resolution mode) with a resolution ∼ 30,000. Mass calibration was performed with sodium formate over a mass range of 50-1200 *m/z*. Data was collected over the 30 min chromatographic separation with a scan time of 0.5 sec. Data-independent acquisition of MS/MS (MS^E^) was performed in the transfer region of the instrument using a 45-60 eV collision energy ramp. Leucine enkephalin was continuously infused throughout the run for post-acquisition mass correction.

### Data Analysis

The small molecules version of Skyline was used for analysis of the lipidomics data (29, 31). Transition lists containing fatty acyl tail compositions of 30:0-35:0 were created for PGs (negative mode, [M-H]^-^ adducts) and LysylPGs (positive mode, [M+H]^+^ adducts). Each lipid species in the transition list contained FA fragments ranging from 13 to 20 carbons, including *d_3_* and *d_4_* isotope-labeled species before and after elongation. Waters .raw files were imported directly into Skyline. Lock mass correction was performed using a 0.05 Da (negative mode) or 0.1 Da (positive mode) window. Ion mobility filtering was performed with a drift time width of 0.60 ms. Overlaid extracted ion chromatograms from the MS1 and MS/MS dimensions were used to identify the fatty acyl tails of lipid species based on their matching retention times and drift times. Peak areas for each lipid species, isomer, and fatty acid fragment were integrated to evaluate the differences between strains and growth conditions.

### Fluorescence Anisotropy

The small molecule probe 1,6-Diphenyl-1,3,5-hexatriene (DPH) was used to evaluate membrane fluidity (32). Pellets from overnight cultures were washed with PBS (pH 7.4) twice and redispersed in fresh PBS. Each suspension was adjusted to 2.0 McFarland units (∼6 x 10^8^ CFU/mL) in 2.5-3 mL of PBS and incubated with 2 µM of DPH for 10 min at 37 °C and 200 rpm. Fully prepared suspensions were kept at 37°C using a water bath until analyzed. Anisotropy measurements were taken on a Jobin Yvon FluoroMax-3 spectrofluorometer with polarizers in the L geometry. DPH was excited at 360 nm and the fluorescence intensity monitored at 430 nm. Samples were maintained at 37°C using the temperature-controlled sample compartment of the spectrofluorometer. A DPH-free cell suspension was used for background correction. Measurements were performed on biological triplicates. The anisotropy of DPH was calculated from the differences in the fluorescence intensity detected with the polarizers in the vertical and horizontal orientations, as described elsewhere (32).

### Cell Surface Charge

Bacteria were cultured overnight as above, washed twice in 20 mM MOPS (MilliporeSigma) buffer, and resuspended to an OD_600nm_ of 1.0 with 20 mM MOPS buffer. Cytochrome c (MilliporeSigma) was added into 1 mL of the bacteria suspension to a final concentration of 0.25 mg/mL. Samples were left at room temperature for 10 min for cytochrome c binding to occur, then centrifuged for 10 min at 12,000xg (Thermo Scientific Sorvall Legend Micro 21R). An eight-point calibration curve was prepared from 0 to 0.25 mg/mL cytochrome c in 20 mM MOPS. Supernatants from the bacteria, blanks, and calibration curve points were aliquoted into a flatbottom 96-well microplate and the absorbance at 530 nm was determined (Agilent/Bio-Tek Epoch2). The percentage of unbound cytochrome c was determined from the calibration curve. A decrease in unbound cytochrome c in the supernatant is indicative of a more negatively-charged cell surface.

### Transmission Electron Microscopy

Bacteria from overnight cultures were pelleted, washed, soaked in 0.1 M PBS containing 2% glutaraldehyde for 1 hr. Each sample was washed 3x with PBS and submerged in 4% noble agar. The agar was allowed to set overnight and then cut into sections. Each section was fixed in 1% osmium oxide buffer for 1 hr, rinsed, and dried via ethanol series and moved to a 50% ethanol/propylene oxide (PO) solution, followed by pure PO for 5 min each. Sample polymerization occurred at 60°C overnight in Embed 812. Hardened samples were trimmed using an ultramicrotome to 50-70 nm sheets. Sheets were placed on a copper grid and soaked in uranyl acetate for 30 min, followed by lead citrate for 5 minutes. Images were taken using a JEOL JM1011 transmission electron microscope (TEM). Measurements of cell wall thickness were performed with ImageJ for 5 cells from each strain and a total of 30 measurements were taken for each cell. Data represents the mean and standard deviation of 150 individual measurements.

### Determination of bkd cluster and pdhB expression using RT-qPCR

Total RNA was first isolated from cultures grown to OD_600_ of 0.600 (mid-exponential phase). Bacterial cells were collected by centrifugation and then resuspended in 1.0 ml RNA-Bee (Tel-Test, Inc., Friendwood, TX) and lysed with 0.1 mm silica-zirconium beads in a BioSpec Mini-Beadbeater. The cells were subjected to six cycles of bead beating (60 seconds each with 1 minute on ice in between cycles). Cell-free supernatant was collected by centrifugation and added with 200 ml of 1-bromo-3-chloropropane (BCP) and vortexed vigorously for 1 min and allowed on ice for phase separation. The aqueous phase was carefully aspirated after a centrifugation for 5 min at 4 °C and the total RNA was precipitated using an equal volume of isopropanol and incubation overnight at −20 °C. Total RNA was then pelleted, washed twice with 70% ethanol and resuspended in water. RNA quantities in these samples were quantified using a Nanodrop spectrophotometer (Thermo Fisher Inc.). These RNA samples were treated with DNAse I (Ambion) to remove contaminating DNA and purified using a RNeasy Kit (Qiagen) following the manufacturer’s instructions and used in cDNA synthesis.

The real-time RT-qPCR reactions were carried out using the Bio-Rad CFX96 Touch. Initially, cDNAs were synthesized using 10 μg of DNase treated total RNA in a 40 μL reverse transcription reaction. The reaction mixes included SuperScript III Reverse Transcriptase (Invitrogen) and random hexamers. A negative control reaction was performed with samples that lacked the reverse transcriptase to confirm that there was no genomic DNA still present. Specific gene primers for both dihydrolipoamide dehydrogenase, *lpdA* (LpdA-F and LpdA-R), the first gene in the *bkd* locus, and pyruvate dehydrogenase, *pdhB* (PdhB-F and PdhB-R), were then used in RT-qPCR using the cDNA samples as template using iQ^TM^ supermix (Bio-Rad) in a two-step PCR. The PCR ran for 40 cycles (95 °C for 30 sec, 60 °C for 15 sec) with an initial denaturation step of 95 °C for 90 sec. In these assays, the transcript levels were normalized to the transcript level for a house keeping gene encoding DNA gyrase (*gyrA*) as the expression of this gene is suggested to remain constitutive (33). The expression levels of *lpdA* and *pdhB* were calculated using the formula 2^(-ΔΔCq)^ (34).

### Creation of Complemented Strains

For genetic complementation with the *pgsA* gene, a 1391 bp DNA fragment was PCR amplified using primers PgsA-F (5’-GGTACCTCGGTGAAGCGCTAAAAGGT-3’) and PgsA-R (5’-TCTAGAAGCGCATGACGTACACTTGA-3’), and *S. aureus* N315 genomic DNA as template. The amplicon represents a fragment starting 416 bp upstream of the *pgsA* start codon and 396 bp downstream of the *pgsA* stop codon that was cloned to the *Kpn*I and *Xba*I sites of a shuttle plasmid pCU1 and subsequently transferred to the *S. aureus* N315-D8 strain (35).

### Daptomycin Survival Assay

N315-D8 (pCU1) and N315-D8 (pCU1-*pgsA*) were cultured overnight in TSB containing 25 µg/mL of chloramphenicol. Pellets from overnight growth were washed and resuspended in TSB containing 50 mg/L CaCl_2_ and 160 µg/mL daptomycin (GoldBio). Cultures were incubated at 37°C with shaking for 4 hrs. Aliquots of the culture were taken at 0 and 4 hrs for colony enumeration. Survival was calculated as the ratio of colonies at 4 hrs versus 0 hrs. The experiment was repeated on three separate days.

### Minimum Inhibitory Concentration Assay

Individual colonies were selected from agar plates and used to prepare 0.5 McFarland suspension in sterile saline. Mueller-Hinton broth supplemented with 30 mg/L CaCl_2_ was inoculated with the bacteria suspension at a dilution factor of 1:100. The inoculated broth was added in a 1:4 dilution into a 96-well plate containing a concentration gradient of daptomycin (0.125 to 64 µg/mL) in Ca^2+^ adjusted MHB. The plate was incubated at 37°C without shaking and MICs interpreted at 24 hrs.

### Data Availability

Raw data files from the RPLC-IM-MS lipidomics analyses are publicly available on the MassIVE Repository (MSV000091690, doi:10.25345/C51Z4235K). The Skyline libraries are available on the PanoramaWeb dashboard for the Hines Lab UGA.

## RESULTS

### BCFA:SCFA Distribution in Membrane Lipids of N315-D8

As previously reported for N315 and N315-D8 (28), we detected a general decrease in the abundance of PG lipid species in strain N315-D8 relative to its isogenic parent strain due to a SNP in *pgsA*. This decrease was not uniform across all lipid species within the PG class, as the lipids with two odd-carbon fatty acyl tails were decreased to a lesser extent than those containing two even-carbon fatty acyl tails or mixed acyl compositions. Using a newly reported RPLC method, we have revealed major differences in the distribution of branched- and straight-chain fatty acyl tails in the membrane lipids of *S. aureus* strain N315-D8 with high-level daptomycin resistance. Representative data from the chromatographic separation of PG isomers arising from different combinations of straight- and branched-chain fatty acyl tails is presented in **Figure 1**. The parent strain, N315, contained PG species with fully branched fatty acyl tails (*b/b*), a mixture of branched and straight chain tails (*s/b*), and a small amount of fully straight acyl tails (*s/s*), as shown for PG 32:0 and PG 33:0 in **Figure 1A,B** (chromatograms for all other PGs can be found in **SI Figure S2**). For N315-D8, the even carbon PGs almost exclusively consisted of fully BCFA tails (**Figure 1A**) while odd carbon PGs were largely mixed SCFA/BCFA (*s/b*) tails with a small percentage of the *b/b* isomer (**Figure 1B**). Collectively, the fatty acyl content of PGs was 83% branched in N315-D8 whereas N315 PGs contained only 64% BCFA (**Figure 2A**). The exact fatty acyl tail compositions of PGs from N315 and N315-D8 can be found in **SI Figures S3 and S4**.

**Figure 1.**
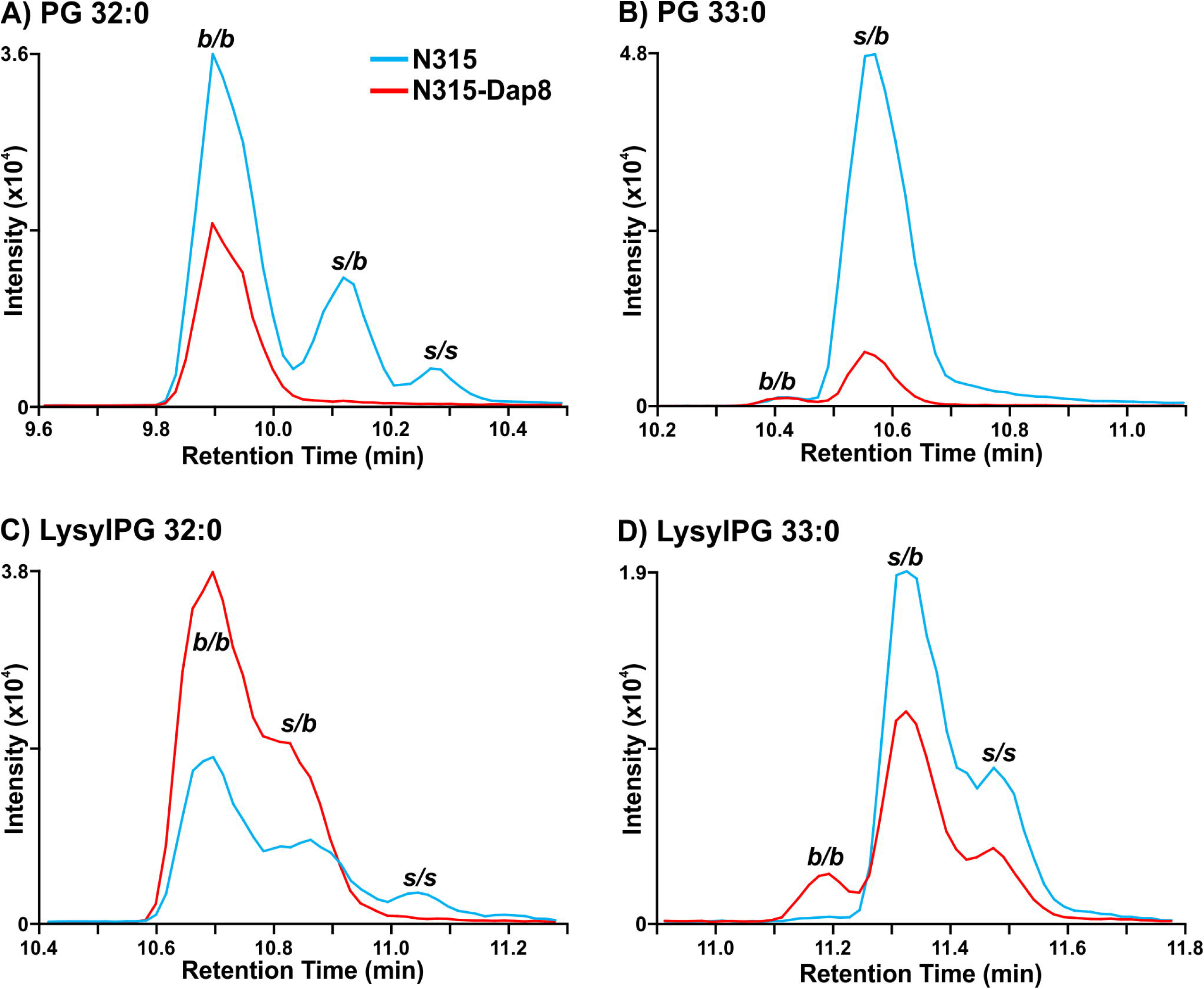
Extracted ion chromatograms of A) PG 32:0, B) PG 33:0, C) LysylPG 32:0, and D) LysylPG 33:0 differences in the distribution of branched- and straight-chain fatty acids between *S. aureus* N315 (blue) and N315-D8 (red) display. *b/b*, two BCFAs; *s/b*, one SCFA and one BCFA; *s/s*, two SCFAs.

**Figure 2.**
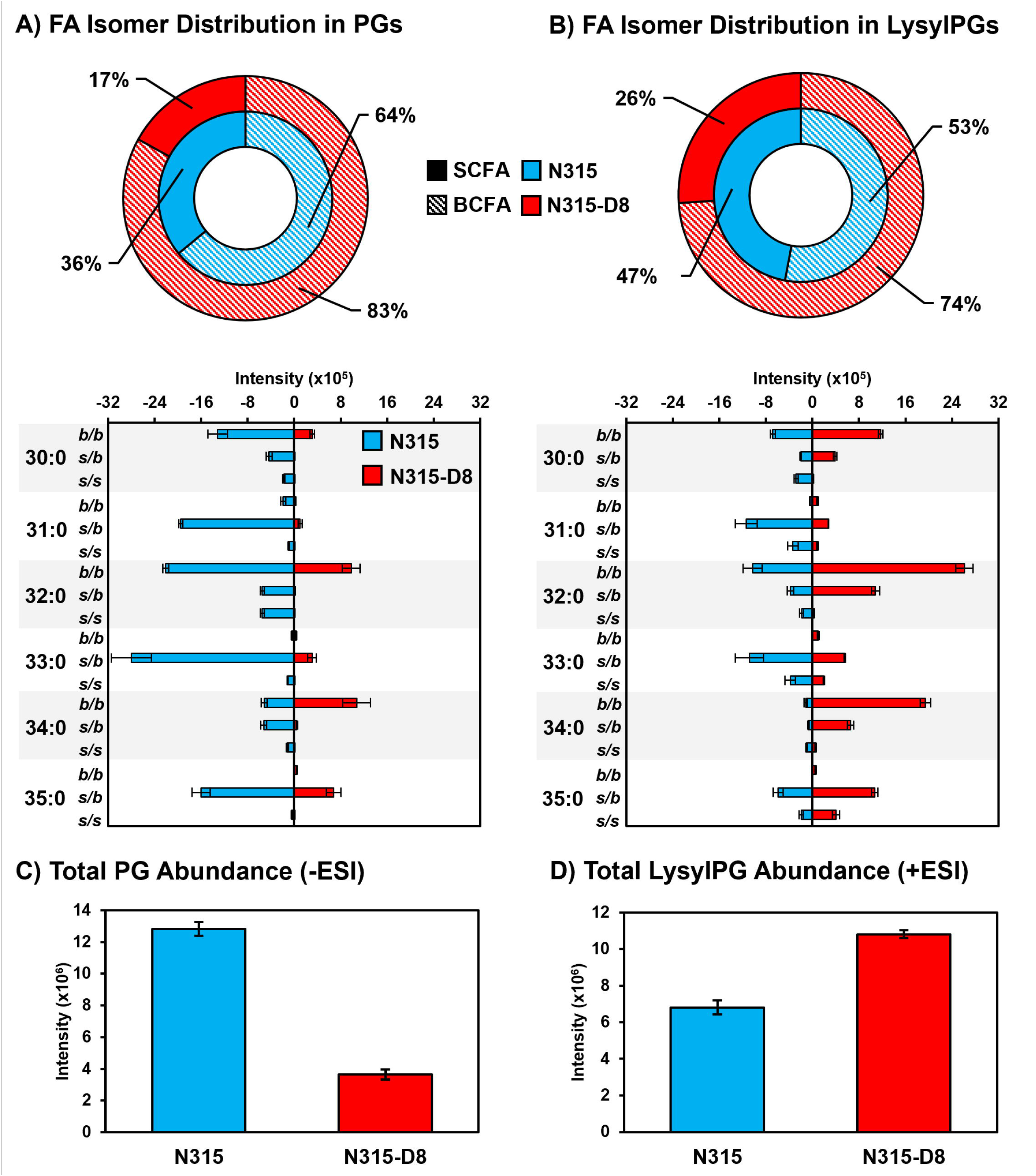
Summary of BCFA and SCFA isomer distributions in the A) PGs and B) LysylPGs for *S. aureus* N315 (blue) and N315-D8 (red). Total levels (all isomers) of C) PGs and D) LysylPGs in N315 and N315-D8. Note that PG and LysylPG abundances are determined from different datasets (negative mode ESI vs. positive mode ESI), and cannot be compared directly due to different ionization efficiencies.

The other major membrane lipids, diglycosyldiacylglycerols (DGDGs) and phosphatidic acids (PAs), followed the same pattern of fatty acyl tail isomers observed for PGs in both N315 and N315-D8 (**SI Figure S5**). However, LysylPGs (**Figure 1C,D**) were elevated in N315-D8 in a manner that varied with the fatty acyl composition and displayed a difference in the distribution of lipid species between the acyl chain isomers. The LysylPGs with total fatty acyl compositions of 30:0, 32:0, 34:0, and 35:0 were elevated in N315-D8 relative to N315 (**Figure 2B**). The elevated LysylPGs were attributed to increases in the *b/b* and *s/b* isomers, whereas only LysylPG 35:0 contained a greater proportion of *s/s* isomer than the N315 parent strain. Although the LysylPGs of N315-D8 contained significantly more BCFAs than those of N315, both strains contained ∼10% more SCFA in LysylPGs than in PGs (**Figure 2B**). Consistent with previous data and the reported mutations in *pgsA* and *mprF*, N315-D8 contained substantially less PGs (**Figure 2C**) and more LysylPGs (**Figure 2D**) compared to the parent N315 strain.

### Characterization of Cell Envelope Properties

The ratio of branched-to-straight chain fatty acids is used as an indicator of membrane fluidity for *S. aureus*, with greater proportions of BCFAs resulting in increased membrane fluidity. The large shift in BCFA:SCFA towards more branched membrane lipids suggests that the membrane of N315-D8 has greater fluidity than N315. The fluidity of a membrane can be evaluated indirectly by measuring the fluorescence polarization, or anisotropy, of DPH. Small decreases in anisotropy values represent a significant increase in membrane fluidity. Fluorescence polarization of DPH in the membranes of N315 and N315-D8 yielded anisotropy values that differed by 17% (0.32 vs 0.27 *a.u.*, respectively) (**Figure 3A**), confirming that the membrane of N315-D8 was more fluid than that of N315.

**Figure 3.**
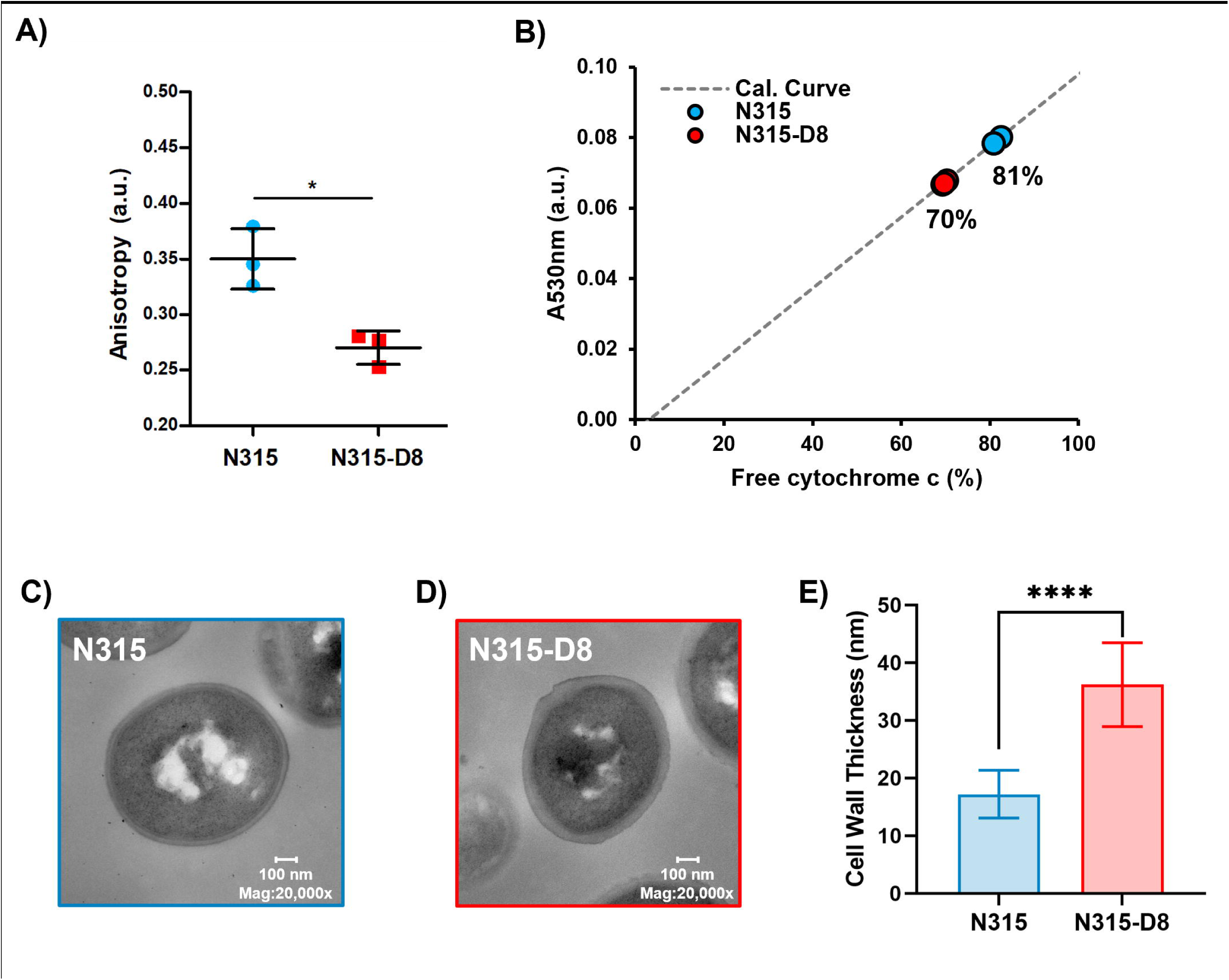
Characterization of A) membrane fluidity for N=3 biological replaces, B) cell surface charge for N=3 biological replicates, and C)-E) cell wall thickness for *S. aureus* N315 and N315-D8. *, *P* ≤ 0.05; **, *P* ≤ 0.01; ****, *P* ≤ 0.0001

The cytochrome c binding assay was used as an indirect assessment of the surface charges for N315 and N315-D8. Using biological triplicates, it was determined that the suspension of N315 left 81.4 ± 0.8% of the cytochrome c unbound whereas as a smaller percentage of cytochrome c (69.7 ± 0.4%) remained unbound in the suspension of N315-D8 (**Figure 3B**). Therefore, it can be concluded that the cell surface of N315-D8 was ∼12% more negatively-charged than that of the parent N315 strain (*P* = 4.8 x 10^-5^). The significant difference in the cell surface charge between N315 and N315-D8 suggested other modification of the cell envelope beyond the cell membrane. TEM measurements of N315 and N315-D8 were performed to evaluate cell morphology and cell wall thickness. N315-D8 had significantly thicker cell walls (**Figure 3C,D**) that were an average of 19 nm thicker than N315 (**Figure 3E**, *P* = 1.7 x 10^-5^).

### Supplementation of N315 and N315-D8 with SCFAs

*S. aureus* is known to utilize exogenous free fatty acids for the synthesis of membrane lipids when they are supplied in the culture medium (36). Based on the disparities in SCFAs noted above, we exploited the FakA/B1 pathway to evaluate the preferences of N315 and N315-D8 for the synthesis of membrane lipids with SCFAs using stable isotope labeled pentadecanoic (SCFA 15:0) and palmitic (SCFA 16:0) acids (**Figure 4**, **Figure 5, and SI Figures S6-S9**). Labeled SCFA usage differed between N315 and N315-D8 with N315 using slightly more *d4*-SCFA 16:0 and significantly less *d3-* SCFA 15:0 than N315-D8 (**Figure 4A and 5A**). Both conditions reduced total BCFA in PGs for N315-D8 (**Figure 4B and 5B**); however, *d3-*SCFA 15:0 incorporated far more than *d4*-SCFA 16:0, resulting in a similar BCFA:SCFA ratio to N315. We observed that *d3-*SCFA 15:0 and *d4*-SCFA 16:0, and their FASII elongation products *d3*-SCFA 17:0 and *d4*-SCFA 18:0, respectively, were incorporated primarily into the *s/b* PGs isomers where they were paired with endogenous BCFAs. Based on the abundance of fatty acyl fragments in the tandem MS data, the exogenous isotope labeled SCFAs were incorporated predominantly onto the *sn-1* position of the glycerol backbone whereas the *sn-2* position was occupied by the endogenous BCFA (**SI Figure S10**). Additionally, we detected a small amount of lipid species having two exogenous isotope-labeled fatty acyl tails (such as PG *d3*-SCFA 17:0/*d3*-SCFA 15:0 in **Figure 4 D,E, and SI Figure S11**). The dual-incorporation events at the *sn-1* and *sn-2* positions were confirmed by the mass shift induced by the isotope labels (*i.e.*, +6 Da and +8 Da relative to endogenous lipids), their elution in the third chromatographic peak ascribed to the *s/s* isomer, and the exclusive presence of isotope-labeled fatty acyl fragments in the tandem MS data. Incorporation of exogenous SCFAs did not increase total PG abundance (*e.g.*, labeled and unlabeled PG species) (**Figure 4C and 5C**) and, despite the clear preference for BCFAs in its endogenous lipid composition, the individual PG species that incorporated deuterated SCFAs were similar between N315-D8 and N315 (**Figure 4D, 4E, 5D and 5E**).

**Figure 4.**
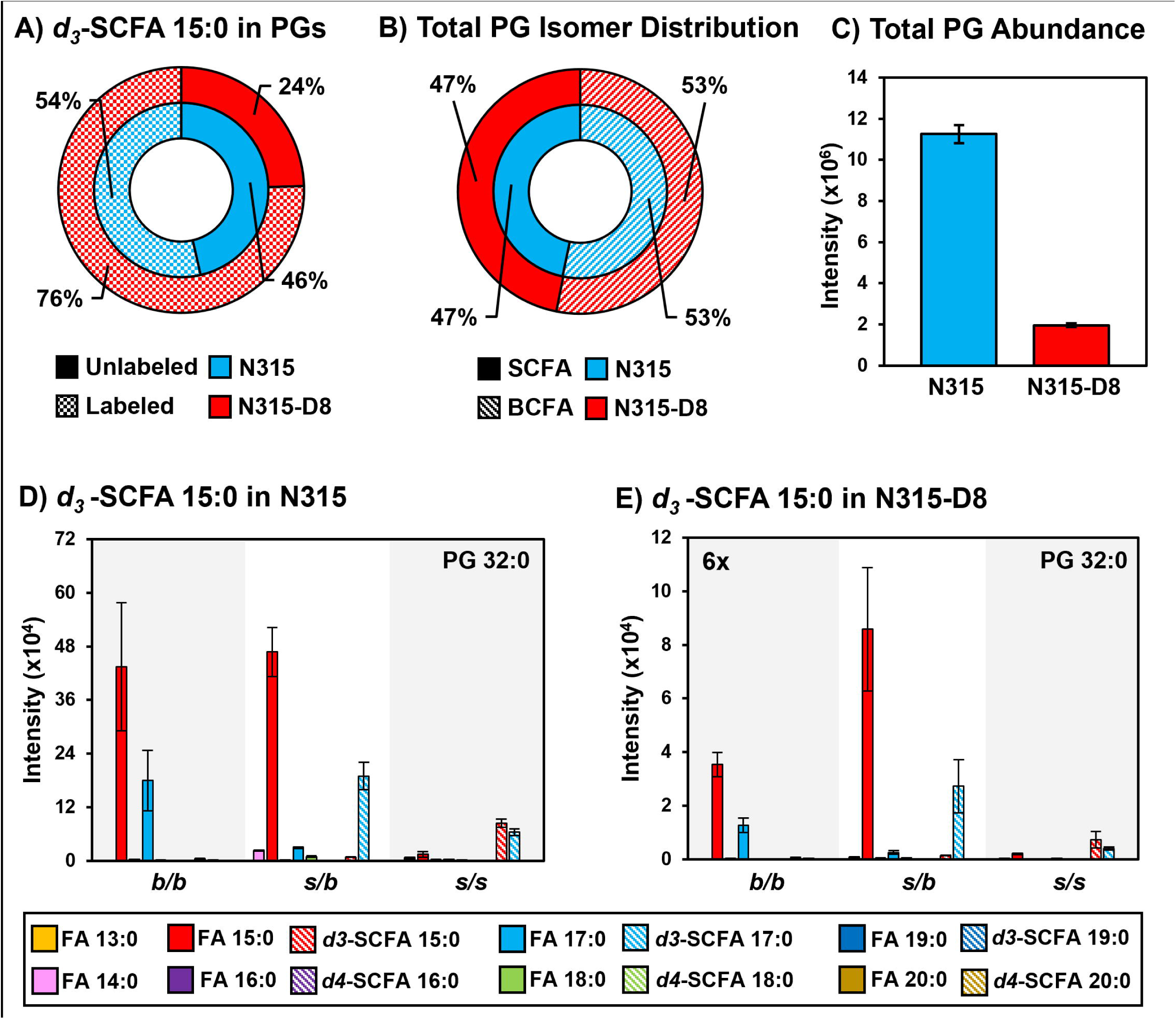
A) Percentage of PGs containing *d3*-SCFA 15:0 versus unlabeled PGs in N315 and N315-D8. B) BCFA and SCFA ratio after supplementation with *d3*-SCFA 15:0. C) Total PG abundance in N315 and N315-D8, including both labeled and unlabeled species. PG 32:0 labeled and unlabeled FA distribution in D) N315 and E) N315-D8.

**Figure 5.**
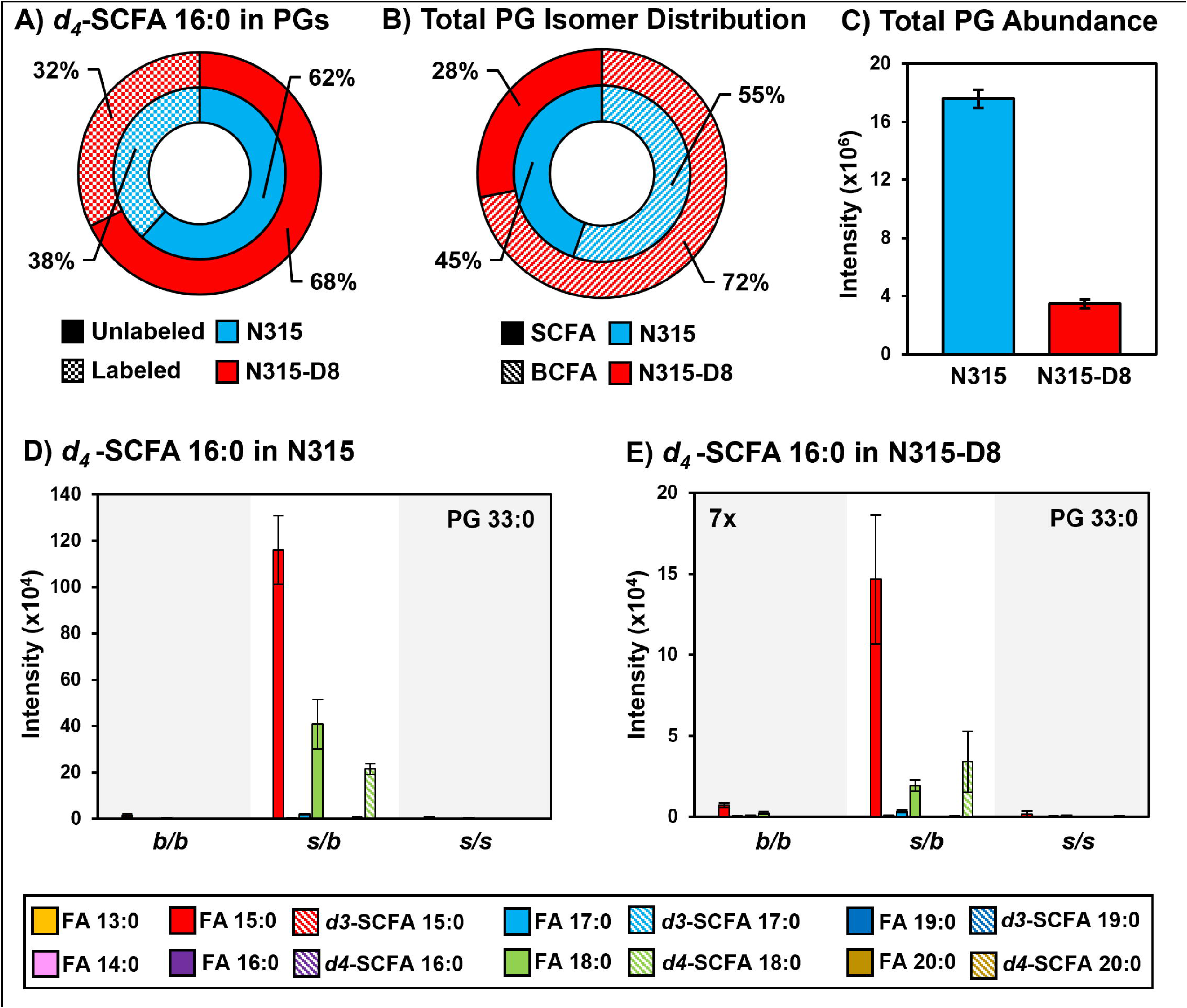
A) Percentage of PGs containing *d4*-SCFA 16:0 versus unlabeled PGs in N315 and N315-D8. B) BCFA and SCFA ratio after supplementation with *d4*-SCFA 16:0. C) Total PG abundance in N315 and N315-D8, including both labeled and unlabeled species. PG 33:0 labeled and unlabeled FA distribution in D) N315 and E) N315-D8.

The incorporation of labeled-SCFAs into LysylPGs was also investigated. As with PGs, the labeled SCFAs were predominantly incorporated into the *s/b* isomers of LysylPGs in N315 and N315-D8 and resulted in a decrease in the amount of *b/b* isomers as endogenous BCFAs were redistributed. However, only trace amounts of doubly-labeled LysylPGs were detected despite the reasonable abundance of their corresponding PG precursors in both N315 and N315-D8 (**SI Figure S12**). These differences in labeled-SCFA utilization between PGs and LysylPGs suggests some regulation or restriction in the pool of PGs that are available for the synthesis of LysylPGs.

Although the balance of BCFA:SCFA was partially restored in N315-D8, the incorporation of exogenous SCFAs did not restore all phenotypes of N315-D8 towards its parent strain. TEM images show that N315-D8 treated with SCFA 15:0 had even thicker cell walls with irregular boarder and distorted cell shapes (**SI Figure 13**). Anisotropy measurements taken on cells grown under the same conditions indicated that N315-D8 was more rigid than its parent strain (**SI Figure 14**; anisotropy *ca.* 15% higher in N315-D8 than N315). However, it is difficult to interpret the anisotropy results in the context of the membrane given the clear distortion of the cell envelope from the TEM analysis.

### RT-qPCR of Genes Involved in BCFA Synthesis

In *S. aureus,* the synthesis of BCFAs involves the formation of short branched-chain acyl-CoAs through the branched-chain α-keto acid dehydrogenase (BKD) complex (37). However, other enzymes are known to influence BCFA synthesis, including pyruvate dehydrogenase (PDH). While inactivation of BKD suppresses BCFA levels, inactivation of PDH results in very high levels of BCFAs (37, 38). The N315-D8 strain not only matched the high BCFA-levels seen in PDH-deficient strains, but also demonstrated the delayed growth (**SI Figure 1**) phenotype of PDH deficiency (38). It was thus speculated that N315-D8 probably lacked or was deficient in PDH activity. To investigate this, the transcription level of the PDH gene cluster (based on *pdhB*) was measured in both N315 and N315-D8 at mid-exponential phase. The data demonstrated that N315-D8 indeed had a very depressed level of expression (20.65 ± 7.13% relative to N315) of the PDH gene cluster. Lipid analysis of mid-exponential phase cultures confirmed that N315-D8 displays a higher BCFA:SCFA ratio than N315 at both timepoints, although the magnitude of the difference is higher at mid-exponential phase (**SI Figure S15**) than stationary phase. Expression levels of the BKD genes, as monitored by the first gene in the cluster, *lpdA*, was unaltered (86.09 ± 2.97% relative to N315). This emphasizes that a PDH-deficiency contributes to slower growth but is conducive for the BCFA production, it was reasoned that a high-level complementation of N315-D8 with PDH-encoding genes can expedite and correct the growth defect in these bacteria. However, when the N315-D8 cells were complemented with the entire PDH gene cluster on a high copy plasmid, the growth of the complemented strain was still very slow compared to the wild-type N315 and the BCFA:SCFA ratio was unaffected (**SI Figure S16**). It is possible that the additional mutations observed in strain N315-D8 are responsible for slowing its growth and complementation with PDH alone is not enough for restoring the growth defect. Another alternative reasoning may be that the proteins that regulate the expression of PDH gene cluster is able to depress the expression of this cluster from the plasmid as well and as a result the N315-D8 strain remains low on PDH activity.

### Complementation of PgsA Mutation Reverses Dap-R Phenotypes and Improves Susceptibility

The lipid composition, total amount of PGs, and cell wall thickness were investigated for strains of N315 and N315-D8 containing the empty pCU1 plasmid and N315-D8 containing a complementary sequence of *pgsA* in pCU1 (**Figure 6**). Complementation of the mutated *pgsA* sequence in N315-D8 reversed the BCFA:SCFA ratio towards the distribution observed in N315 and N315 (pCU1) (**Figure 6A**). Although SCFA supplementation also modified the BCFA:SCFA ratio in N315-D8, the distribution of PG species upon SCFA 15:0 supplementation was shifted towards PG 34:0 (containing 15:0 and 19:0 FAs, **SI Figure S17**) rather than evenly distrusted across PGs as seen in **Figure 6A**. The total amount of PGs in N315-D8 (pCU1-*pgsA*) was elevated 2-fold compared to N315-D8 with the empty pCU1 plasmid (**Figure 6B**). However, the complementation did not fully restore PG levels to the amount present in N315 (pCU1). Levels of both DGDGs and LysylPGs were reduced 2-fold upon complementation of the *pgsA* mutation (**SI Figure S18**), and the BCFA:SCFA ratios of both classes more closely matched the distributions observed for N315. The cell wall thickness (**Figure 6C** and **SI Figure S19**) of N315-D8 was reduced upon complementation of the *pgsA* mutation and there was no significant difference between the cell walls of N315 (pCU1) and N315-D8 (pCU1-*pgsA*).

**Figure 6.**
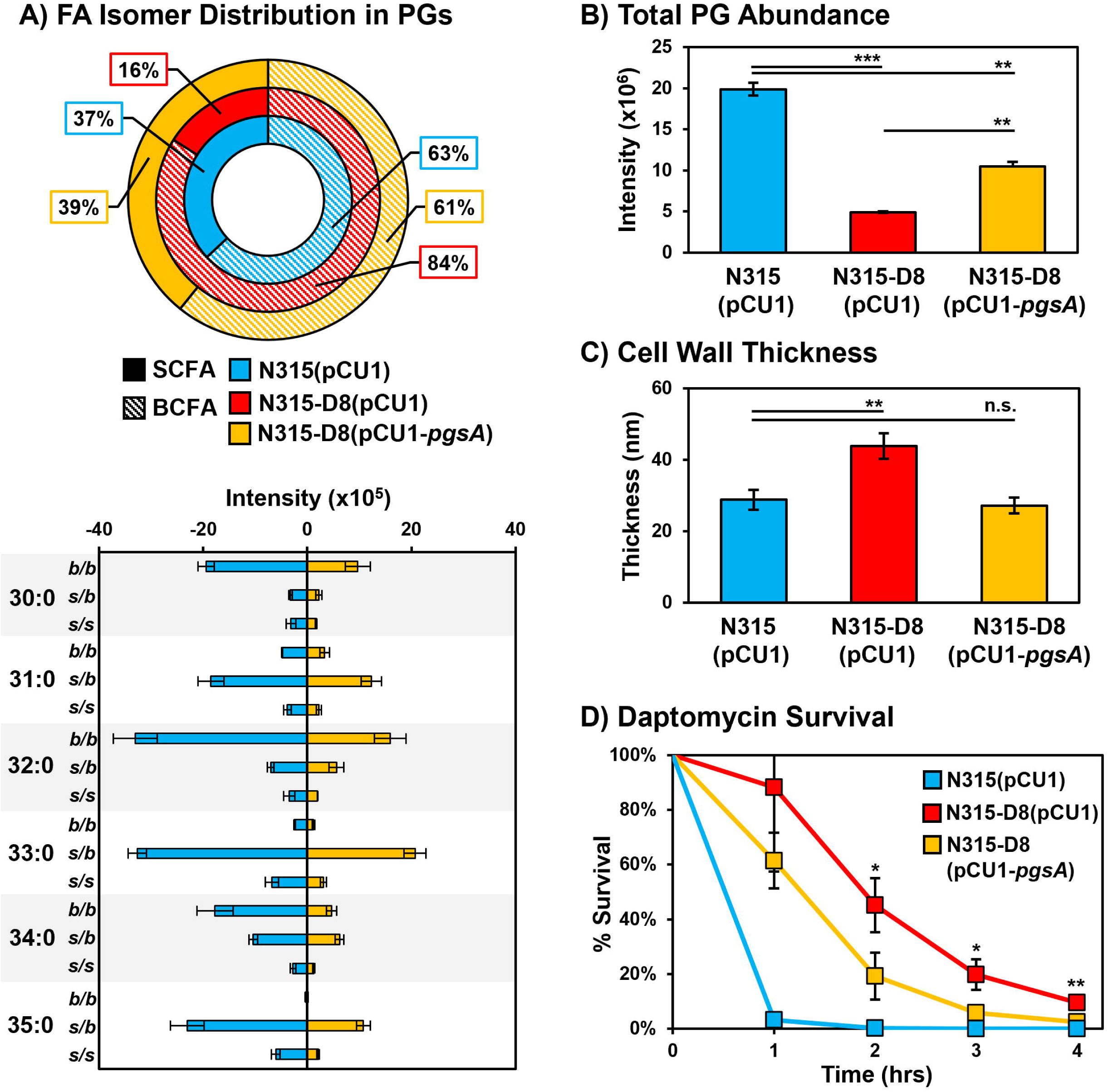
A) Summary of BCFA and SCFA ratios in N315 (pCU1), N315-D8 (pCU1), and N315-D8 (pCU1-*pgsA*). B) Total abundance of PGs in N315 (pCU1), N315-D8 (pCU1), and N315-D8 (pCU1-*pgsA*). C) Cell wall thickness measurements for N315 (pCU1), N315-D8 (pCU1), and N315-D8 (pCU1-*pgsA*). D) Percentage of N315 (pCU1), N315-D8 (pCU1) and N315-D8 (pCU1-*pgsA*) cells that survived a 160 µg/mL daptomycin challenge over 4 hr period. *, P<0.05, **, P<0.01, ***, P<0.001

The partial restoration of PG levels, BCFA:SCFA ratio, and cell wall thickness in N315-D8 (pCU1-*pgsA*) led us to investigate whether the complementation of the *pgsA* mutation was effective in restoring daptomycin susceptibility. N315-D8 (pCU1-*pgsA*) and the empty plasmid control were exposed to 160 µg/mL of daptomycin in TSB supplemented with calcium and the CFU was enumerated every hour for 4 hr. After 2 hr, there was a 26% reduction in CFU for N315-D8 (pCU1-*pgsA*) relative to the strain containing the empty plasmid (**Figure 6D** and **SI Figure S20**). Although the survival of N315-D8 (pCU1-*pgsA*) and N315-D8 (pCU1) normalized towards hr 4, there remained a 7% reduction in CFU for the complement strain relative to N315-D8 containing the empty plasmid. These data suggest that the increase in PG levels coupled with restoration of typical cell wall thickness and membrane fluidity (based on BCFA:SCFA ratio) that were imparted by the functional *pgsA* gene led to an increase in daptomycin susceptibility. We observed that the *pgsA* complement in N315-D8 did in fact result in a 2-fold decrease in the daptomycin minimum inhibitory concentration (**SI Table S2**), whereas the empty plasmid and the attempted *pdh* complement had no effect on daptomycin susceptibilities.

## DISCUSSION

Using high-resolution reversed-phase liquid chromatography and mass spectrometry, the distributions of branched-chain and straight-chain fatty acids were detected within the major lipid classes of *S. aureus*. As PGs are highly-abundant and central to lipid biosynthesis in *S. aureus*, the BCFA:SCFA ratio is often assumed to be similar between PGs and related lipids such as LysylPGs, PAs, and DGDGs. The parent N315 strain investigated here yielded nearly identical BCFA:SCFA distributions between PGs and LysylPGs at 52-53% BCFA and 47-48% SCFA. A 50/50 distribution of BCFAs to SCFAs is typical for *S. aureus*, which has a preference for at least one BCFA in the acyl tails of its membrane lipids, and does not vary through the growth phases based on our data and other measurements of BCFA:SCFA in exponential phase cultures (38–40).

Investigation of the membrane lipids in the high-level daptomycin resistant *S. aureus* strain N315-D8 revealed disparities between the BCFA:SCFA distributions of PGs and LysylPGs in addition to the anticipated alterations to their overall abundances due to *pgsA* and *mprF* mutations. A previous report on the lipid profile of N315-D8 revealed a net decrease in the abundance of PGs, but an acyl tail specific pattern of increased abundance for PGs containing odd carbon, and likely branched-chain, FAs (28). In the present study, we detected a 31% increase in BCFAs within the PGs of N315-D8 compared to its parent strain (83% vs. 52% BCFA) that is consistent with the previous study. Membrane fluidity measurements based on fluorescence anisotropy of the membrane probe DPH confirmed that the bulk membrane of N315-D8 was substantially more fluid than that of N315.

LysylPGs in N315-D8 contained ∼11% more SCFAs than the PGs (26% vs 17% SCFA, respectively). Since PGs are the direct biosynthetic precursor to LysylPGs, the significant difference in the distributions of BCFAs and SCFAs in these two lipid classes suggests that the LysylPGs were synthesized from a specific subset of PGs with slightly more SCFAs than suggested by our bulk lipidomics measurements. These data may represent a local enrichment of SCFA-containing PGs in the vicinity of the lysyltransferase, MprF, that could be driven by a fluidity preference of the protein. MprF is a large membrane protein with 14 transmembrane domains that performs the dual functions of LysylPG synthesis and translocation from the inner to outer membrane leaflet (41, 42). The mutation present in N315-D8, L826F, occurs in the lysinylation domain of the protein that remains on the cytoplasmic side of the membrane and is not membrane bound (41, 42). Although among one of the more common mutations detected in daptomycin-resistant *S. aureus*, the presence of the L826F mutation itself is not enough to confer daptomycin resistance (24). We detected elevated LysylPG abundance in N315-D8 relative to N315, but the cytochrome c binding experiments indicated that N315-D8 in fact had a more negatively-charged membrane than N315. It is possible that L826F MprF produces more LysylPGs in the inner membrane leaflet in the absence of the translocase promoting mutations that more reliably confer daptomycin resistance (24). The substantially thicker cell wall of N315-D8 likely contributed to the increased negative charge of the cell envelope, as well.

The combination of increased BCFAs and reduced total levels of PGs suggest that the fluidity-promoting BCFAs enable the viability of cells in response to the dramatic reduction in total PGs levels caused by the *pgsA* mutation. We initially suspected that the increase in BCFAs was regulated only at the level of fatty acid biosynthesis based on the previously reported *yycG* mutation in N315-D8. The two-component *yycFG* regulatory system is reported to regulate fatty acid biosynthesis in other Gram-positive organisms (43). The site of the mutation, M426I, detected in N315-D8 is within the phospho-acceptor region and may affect the ability of the histidine kinase, *yycG*, to respond to stimuli. In *Streptococcus pneumoniae*, over-expression of unphosphorylated YycF led to increased expression of the genes that encode proteins involved in fatty acid elongation machinery (43). This in turn increased the length of fatty acids synthesized by the organism, which would promote a more ordered, or rigid, membrane. The evidence that pyruvate dehydrogenase, *pdhB*, transcripts are suppressed in N315-D8 and the inability to restore PDH transcription upon complementation are both consistent with an alteration in the regulation of fatty acid synthesis (38), but it is unclear at this time whether or how the *yycG* mutation might contribute to that process.

The *yycFG* system is more commonly known for its role in cell wall metabolism through the activation of autolysin production. A previous investigation of *S. aureus* YycFG suggested that the kinase activity of YycG is suppressed when the fluidity of the local membrane environment is too high due a weakened cell wall (44). This in turn halts the YycF activation of autolysin production so that cell wall integrity can be restored. Daptomycin-resistant isolates of *S. aureus* with mutated *yycG* alone often present with both increased cell wall thickness and increased membrane fluidity (10, 13, 45). However, the thickened cell wall in N315-D8 does not appear to be the result of the strain’s *yycG* mutation. We observed a decrease in the cell wall thickness of our daptomycin-resistant mutant upon complementation of the *pgsA* mutation with the wild-type sequence. Cell wall thickening, in addition to reduced PG levels, has been reported in daptomycin-resistant isolate of *B. subtilis* containing a mutation in *pgsA* (25). There, it was reported that allelic replacement of the *pgsA* mutation with a wild-type copy was able to restore PG amounts but the effect on cell wall thickness was not investigated. Complementation of the *pgsA* mutation not only increased PGs, but also resulted in decreased levels of DGDGs and LysylPGs in N315-D8 (pCU1-*pgsA*). However, failure of the *pgsA* complement to fully restore PG levels suggest that other mutations within N315-D8 (see **SI Table S1**) are contributing to daptomycin resistance, or that PGs are being diverted to other species upon restoration of PgsA function. A potential link between the PGs levels and cell wall is lipoteichoic acid (LTA), which is synthesized from the phosphoglycerol headgroup of PG lipids. In *B. subtilis*, elimination of LTA synthase (LtaS) resulted in a thicker cell wall that caused defects in cell morphogenesis and division (25). It is well documented that LTA plays an essential role in regulating autolysis in *S. aureus*, and strains of *S. aureus* lacking *ltaS* have decreased autolytic activity and thick cell walls (46). Thus, it is plausible that the restoration of *pgsA* in N315-D8(pCU1-*pgsA*) does not fully restore membrane PG levels because PGs are instead diverted to restore normal cell wall thickness through the synthesis of LTA and increased autolysis.

The decreased survival of N315-D8(pCU1-*pgsA*) relative to N315-D8(pCU1) when challenged with daptomycin does suggest that PG levels, cell wall thickness, and BCFA:SCFA distribution are contributing to, but cannot fully explain, the strain’s high-level daptomycin-resistance. Elevated levels of LysylPGs, reduced membrane PGs and cell wall thickness are among the most widely investigated phenotypes of daptomycin resistance (21, 23). From those studies, it is clear that there is more than one route to the development of daptomycin resistance. Although it is not possible to point to a single determinant for daptomycin resistance for N315-D8 given the presence of other mutations (see **SI Table S1**), it is evident that PgsA plays a larger role in *S. aureus* cell envelope homeostasis than phosphatidylglycerol synthesis alone. The observation of altered membrane fluidity is in agreement with newer evidence on daptomycin’s mechanism of action though. At concentrations close to the MIC, *in vivo*, it is believed that the primary effect of daptomycin is on cell wall synthesis, with ion leakage and pore formation being secondary effects (1, 5). Muller et al. provided evidence that daptomycin inserts into regions of increased fluidity in the membrane causing rearrangement of fluid lipid domains and detachment of membrane-associated lipid II synthase MurG and other peripheral membrane proteins (2). In an expansion of this model Grein et al. proposed that Ca^2+^-daptomycin forms a tripartite complex with undecaprenyl-cell wall precursors and phosphatidylglycerol (47). In *S. aureus* the complex is preferentially formed in the septal region divisome complex. In daptomycin-resistant strains such as strain N315-D8 deficient in phosphatidyl glycerol, with increased levels of BCFAs and membrane fluidity, formation of the tripartite complex may be impeded and deleterious delocalization of the peptidoglycan biosynthesis machinery, and membrane rearrangements, may be mitigated by a hyperfluid membrane.

## ACKNOWLEDGEMENTS

This work was supported NIH/NIAID (K22AI143919 to K.M.H; R01AI173144 to K.M.H., B.J.W., and V.K.S.). Additional support to K.M.H. was provided by the University of Georgia Department of Chemistry, Franklin School of Arts and Sciences, Office of Research, and the Office of the Provost. The authors thank Prof. Libin Xu and Prof. Brian Werth of the UW School of Pharmacy for sharing *S. aureus* strain N315-D8. K.M.H. and C.D.F. appreciate the assistance of the Georgia Electron Microscopy Center staff and the Urbauer Lab for providing training on and access to the Jobin Yvon FluoroMax-3 spectrofluorometer.

