## Supporting Information for "Defective pgsA contributes to increased membrane fluidity and cell wall thickening in S. aureus with high-level daptomycin resistance"

***for***

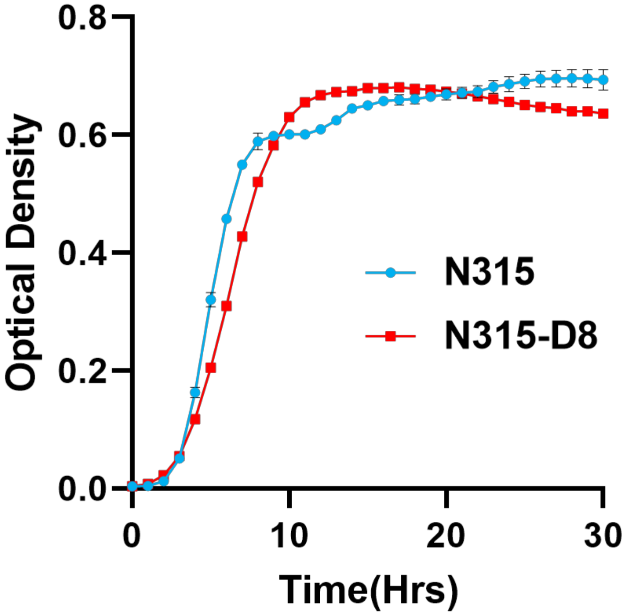


**Figure S1.** Growth curves for N315 and N315-D8 in TSB. Optical density was monitored at 600nm.

**Table S1.** Mutations detected in N315-D8 as previously reported by Hines et al., 2017.

| Predicted Gene Product | Nucleotide Change in N315-D8 | Predicted Amino Acid Change in N315-D8 | Predicted Protein Function |
|---|---|---|---|
| *yycG* | 1278 G → A | M426I | Fatty acid biosynthesis<br>Cell wall biosynthesis |
| *pgsA* | 403 A → G | K135E | PG biosynthesis |
| *mprF* | 2476 C → T | L826F | LysylPG biosynthesis |
| *SA0567* | 865 C → T | Q289* | Iron complex transport |
| *norA* | 737 G → C | G246A | Quinolone resistance protein |
| *rnr* | 1819 G → T | E607* | Ribonuclease |
| *spoIIIE* | Deletion of 2059 C | Q687 frameshift | DNA translocase |

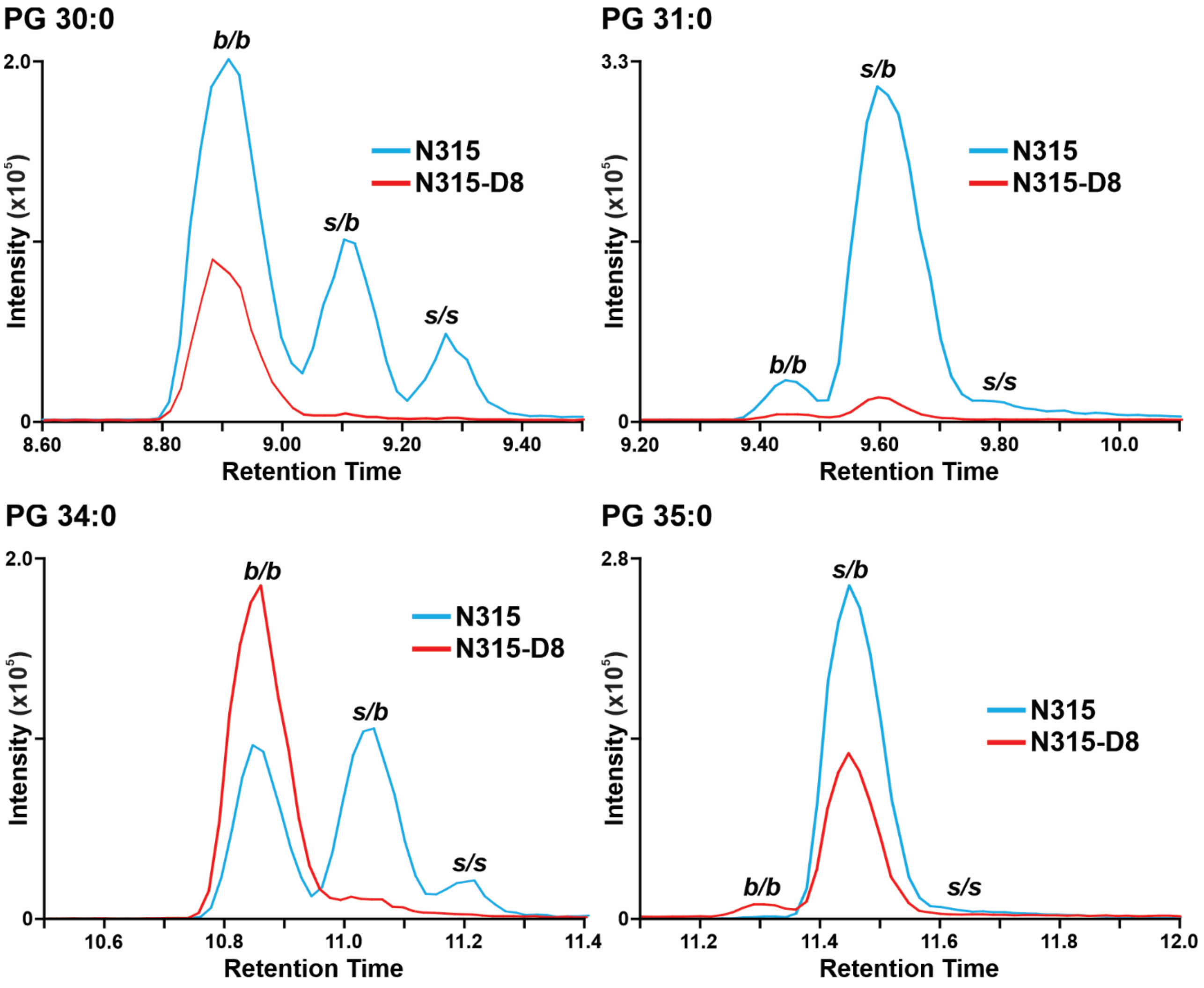


**Figure S2.** Extracted ion chromatograms for PGs in N315 and N315-D8.

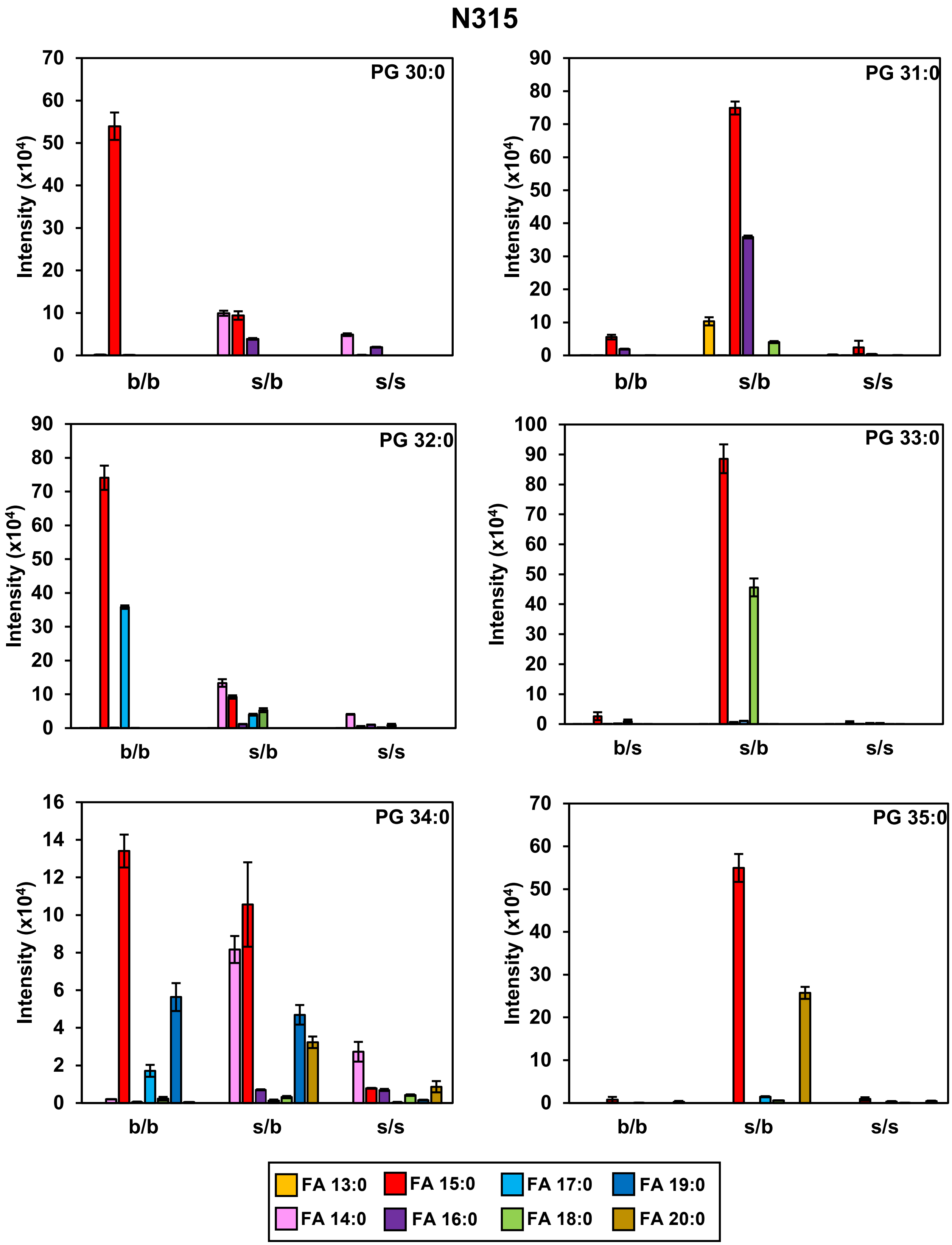


**Figure S3.** Fatty acyl tail composition of PGs in N315 determined by MS/MS.

**N315-D8**

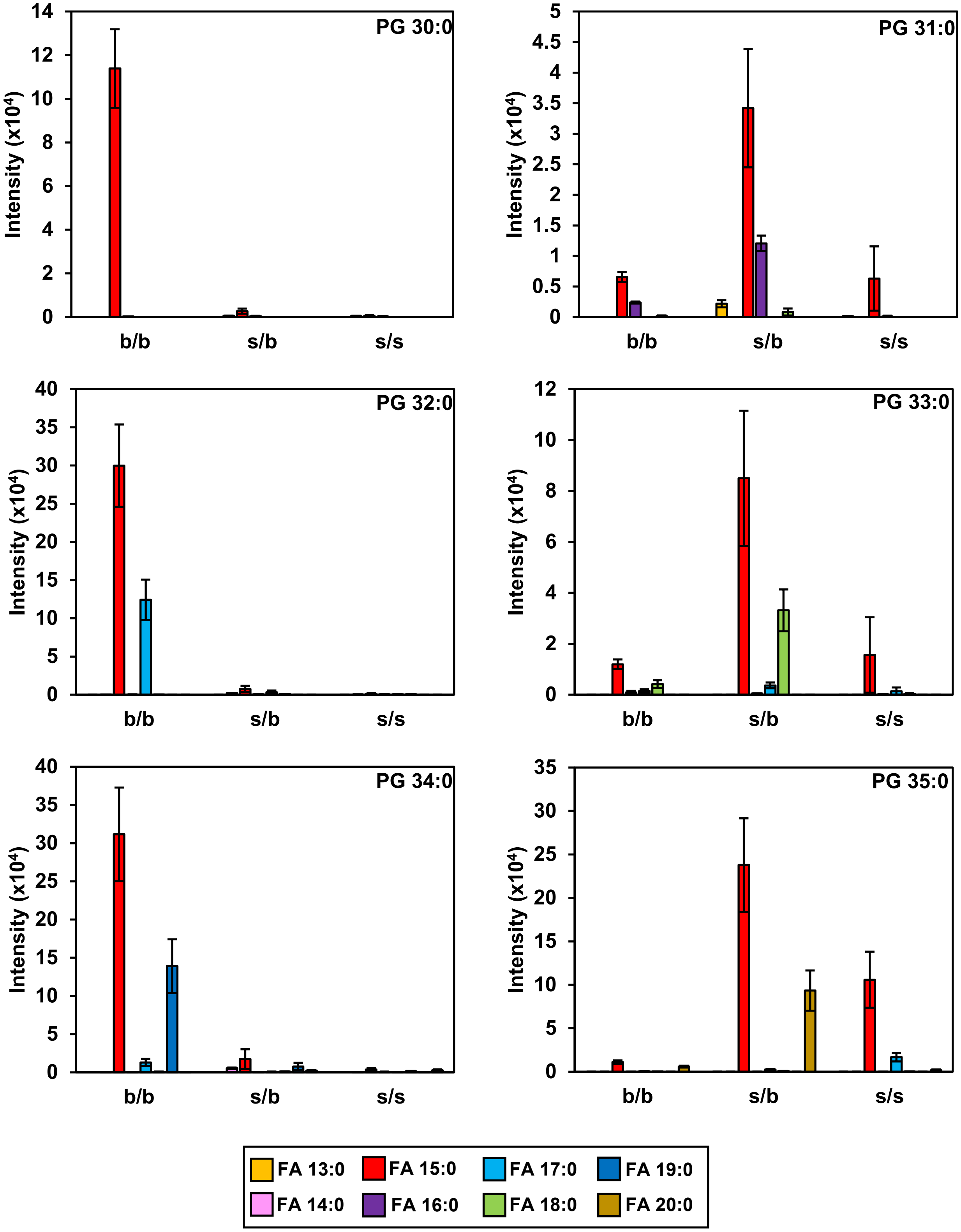


**Figure S4.** Fatty acyl tail composition of PGs in N315-D8 determined by MS/MS.

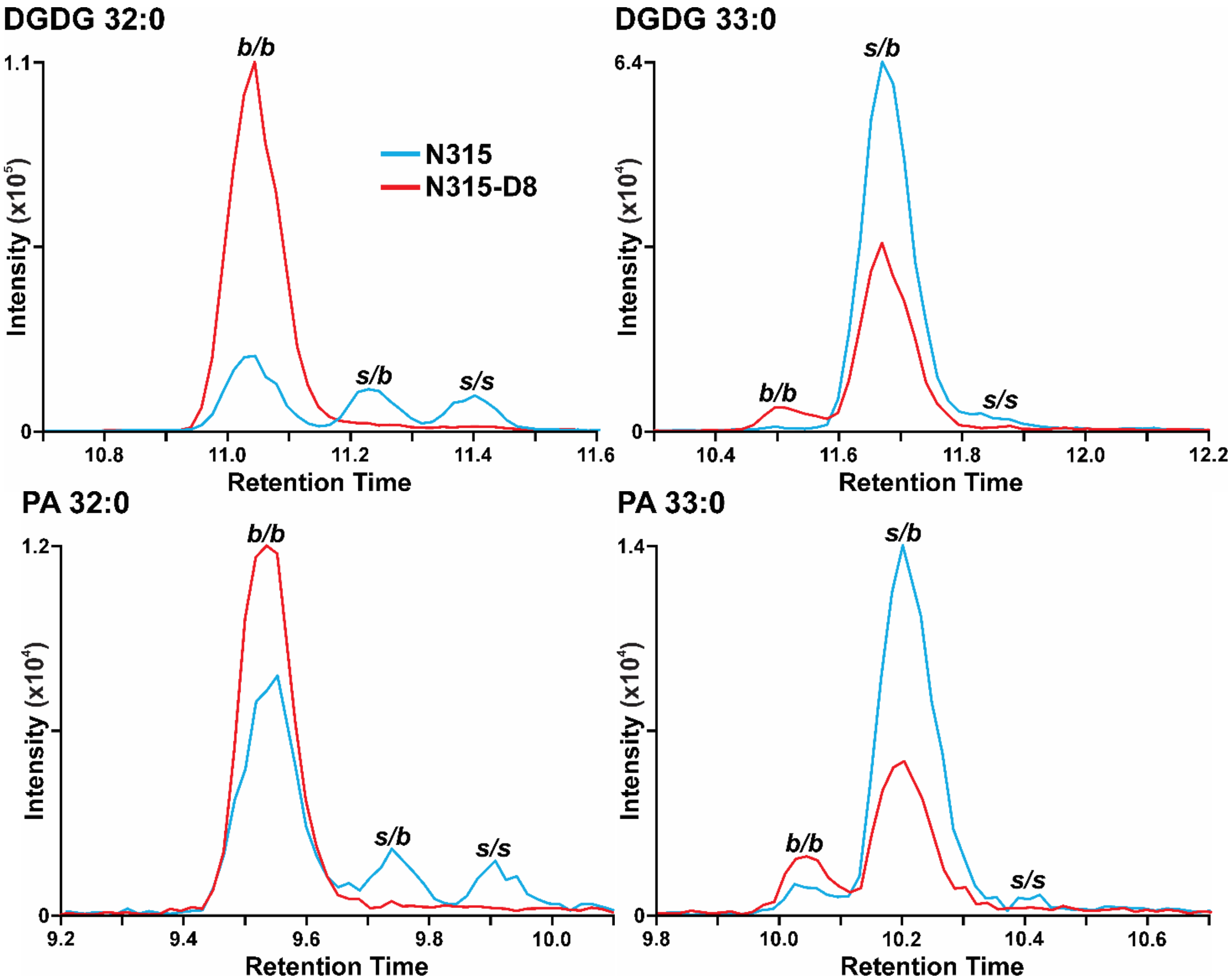


**Figure S5.** Extracted ion chromatograms of A) DGDG 32:0, B) DGDG 33:0, C) PA 32:0, and D) PA 33:0 for N315 (blue) and N315-D8 (red).

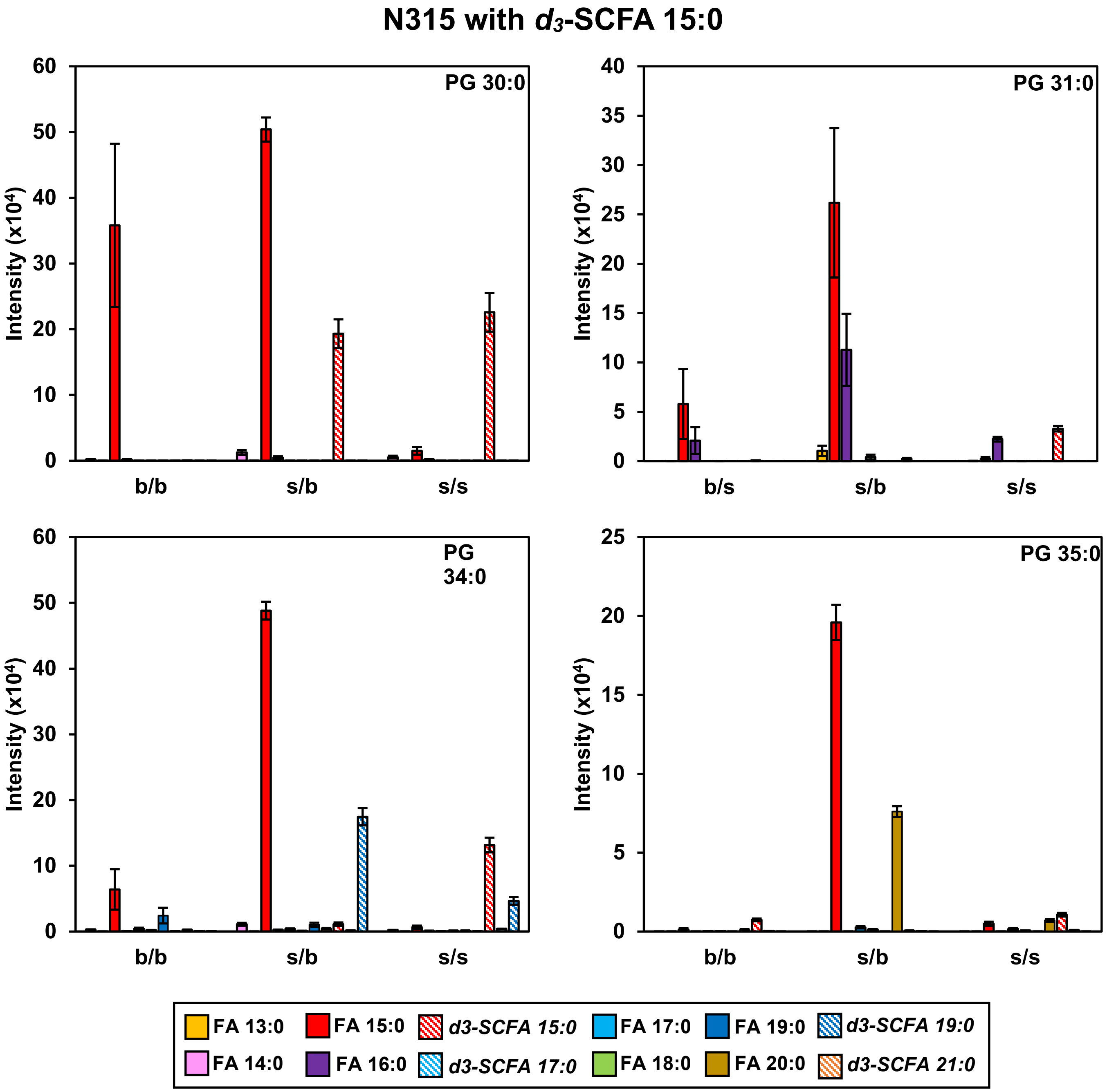


**Figure S6.** Fatty acyl tail compositions of PGs in N315 when grown in TSB containing $d_3$-SCFA 15:0.

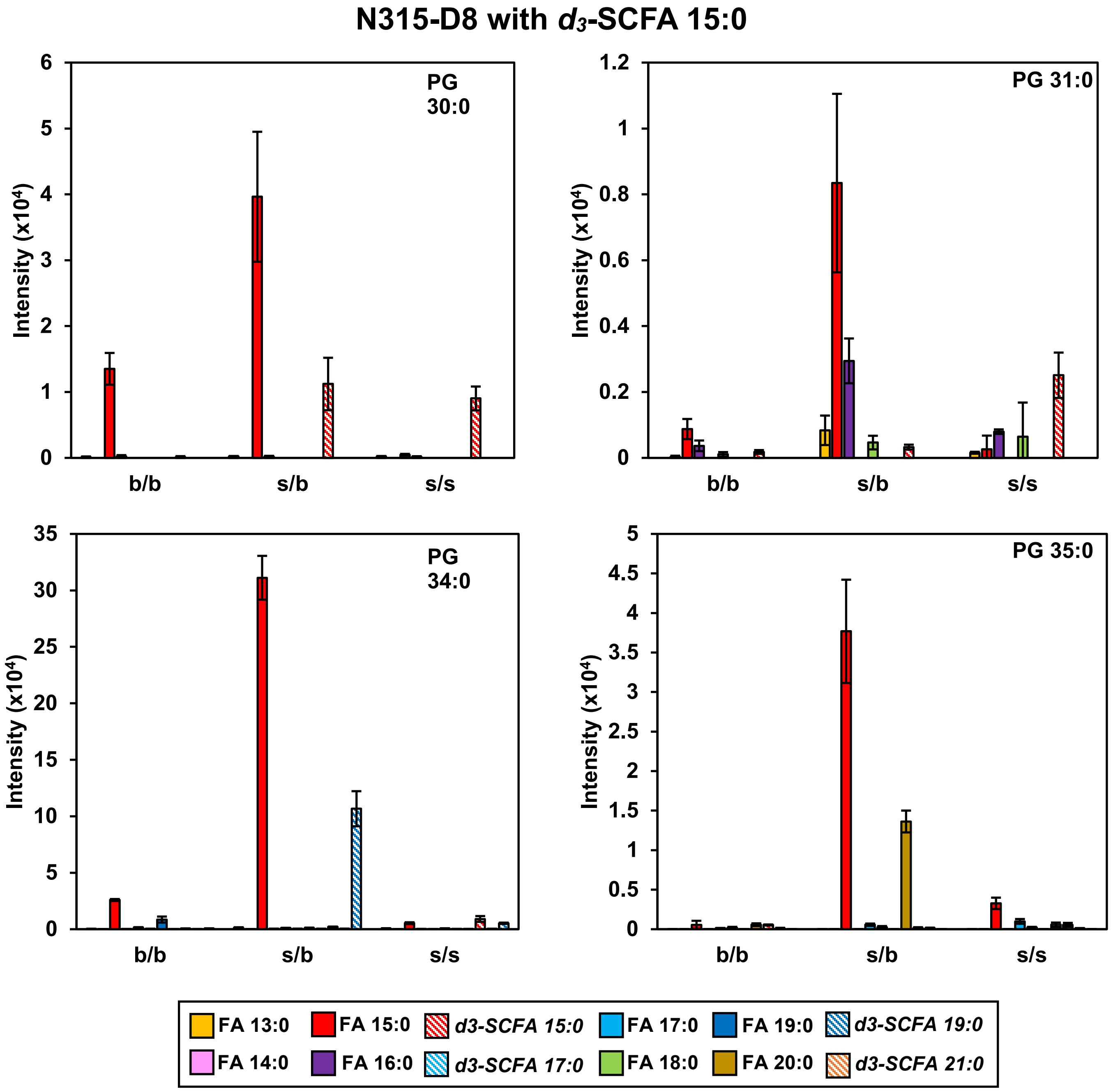


**Figure S7.** Fatty acyl tail compositions of PGs in N315-D8 when grown in TSB containing $d_3$-SCFA 15:0.

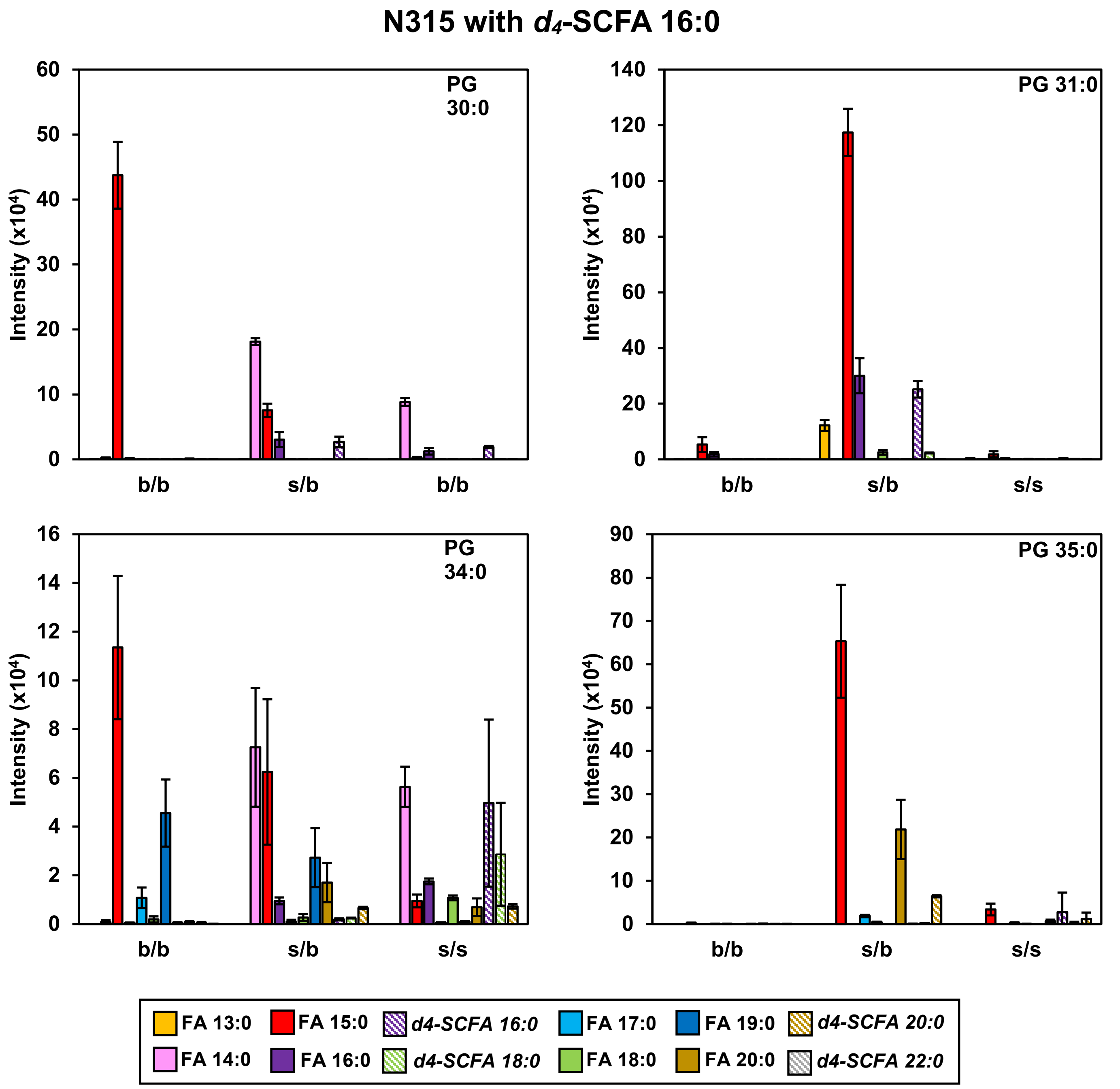


**Figure S8.** Fatty acyl tail compositions of PGs in N315 when grown in TSB supplemented with $d_4$-SCFA 16:0.

### N315-D8 with $d_4$-SCFA 16:0

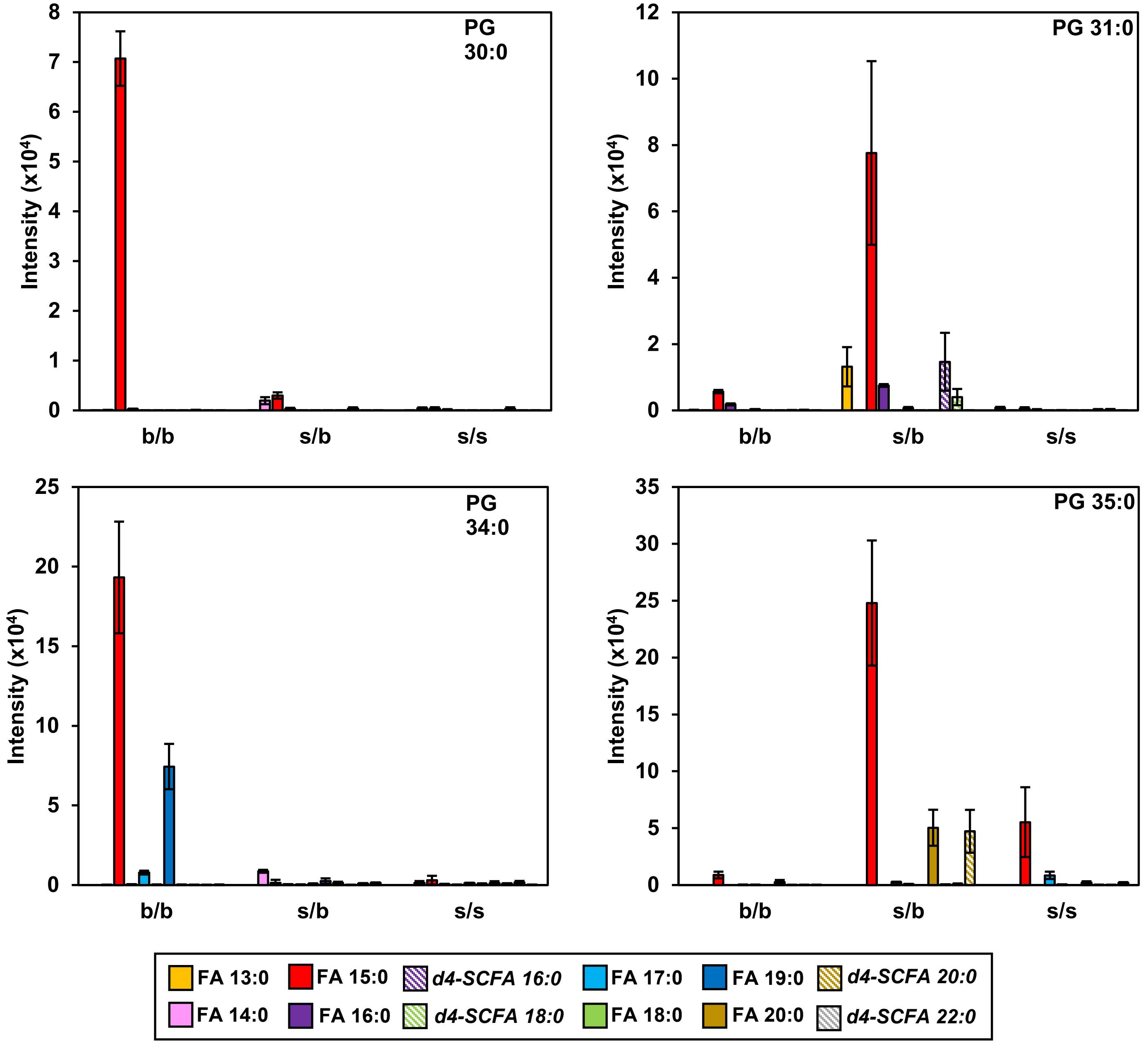


**Figure S9.** Fatty acyl tail compositions of PGs in N315-D8 when grown in TSB supplemented with $d_4$-SCFA 16:0.

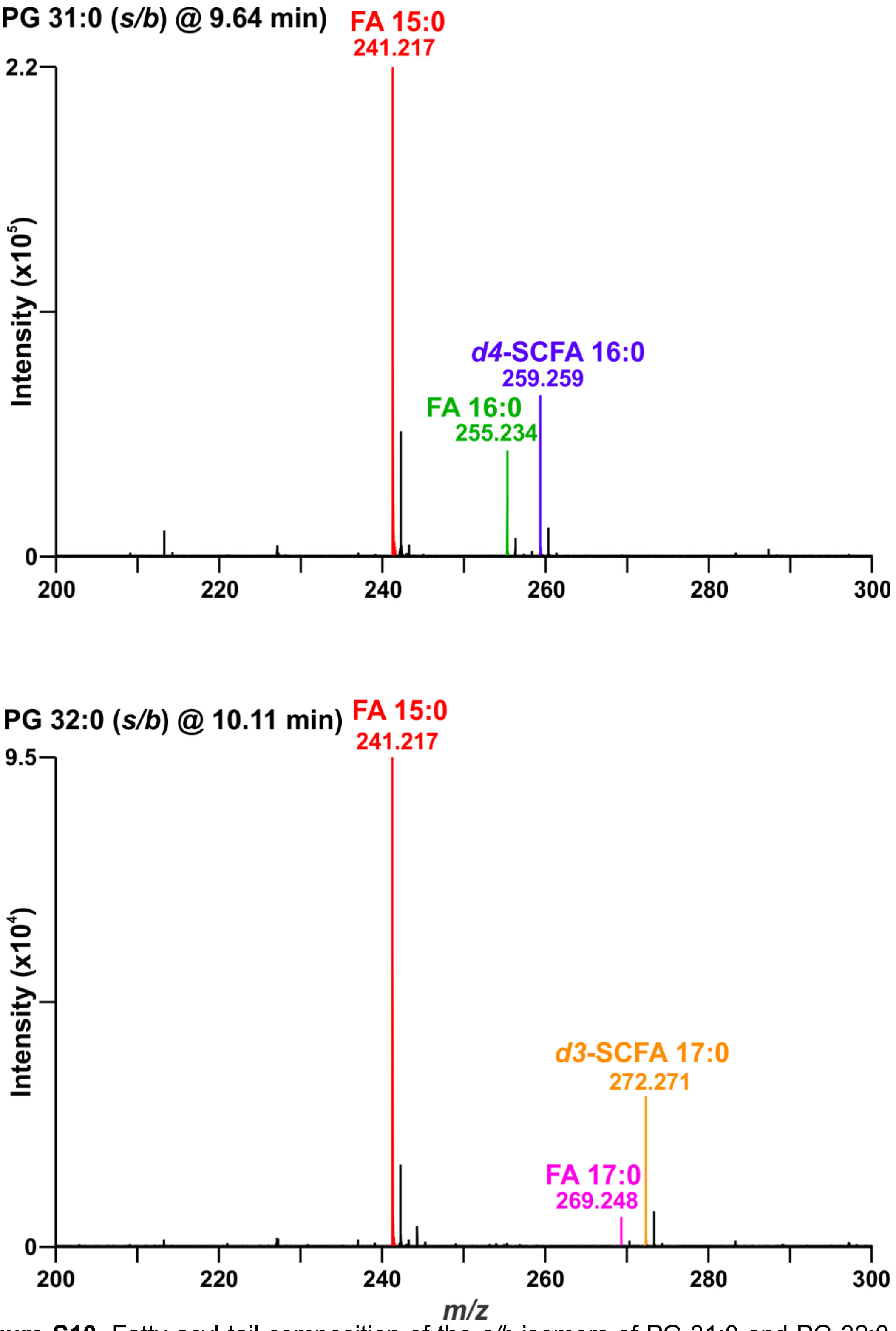


**Figure S10.** Fatty acyl tail composition of the *s/b* isomers of PG 31:0 and PG 32:0 in N315 after supplementation with $d_4$-SCFA 16:0 and $d_3$-SCFA 15:0 show that exogenous FAs incorporate at the *sn-1* position based on acyl tail fragment intensities.

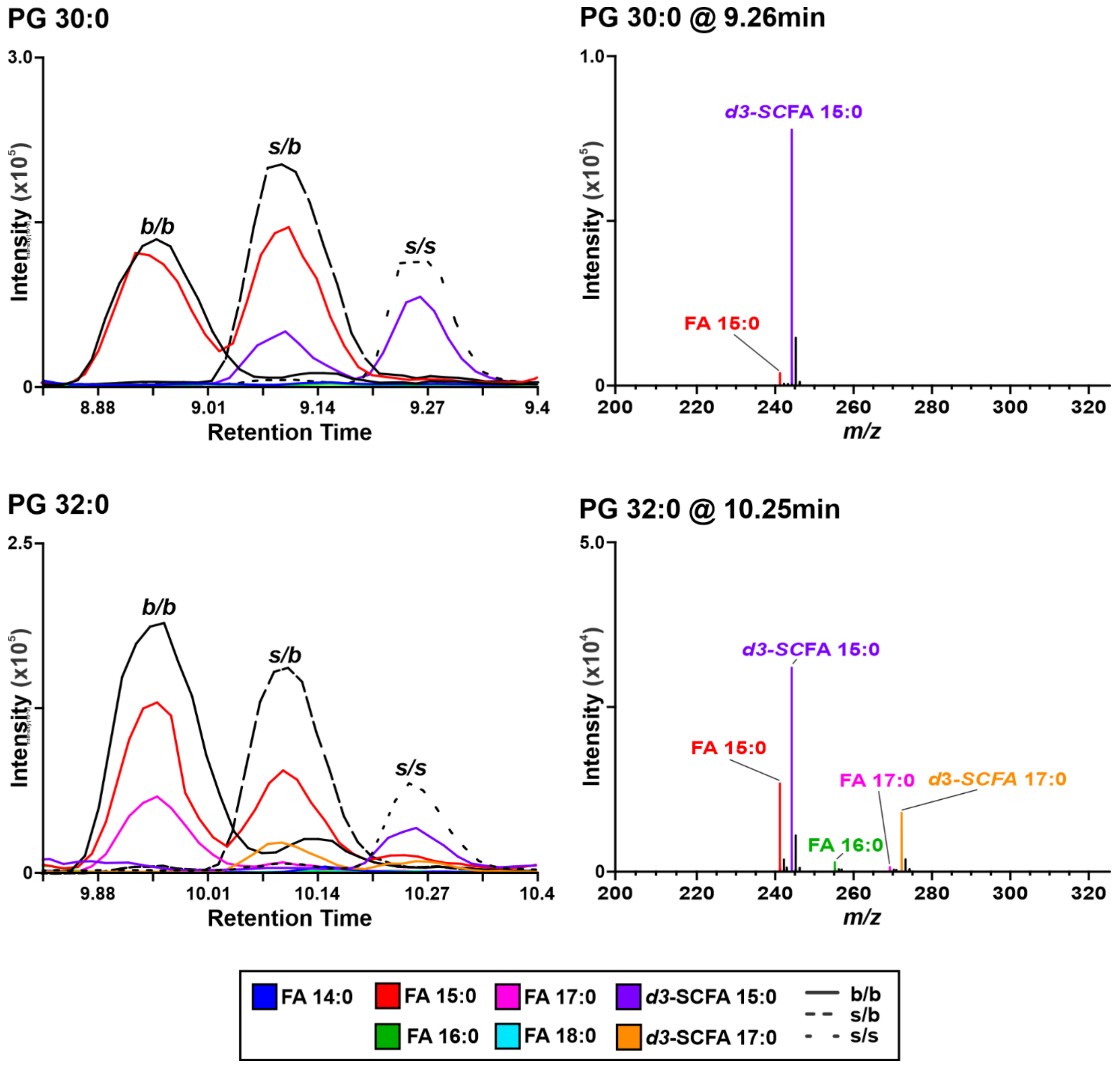


**Figure S11.** Fatty acyl tail composition of the *s/s* isomers of PG 30:0 and PG 32:0 in N315 after supplementation with $d_3$-SCFA 15:0 show the presence of PGs with fully-exogenous acyl tails.

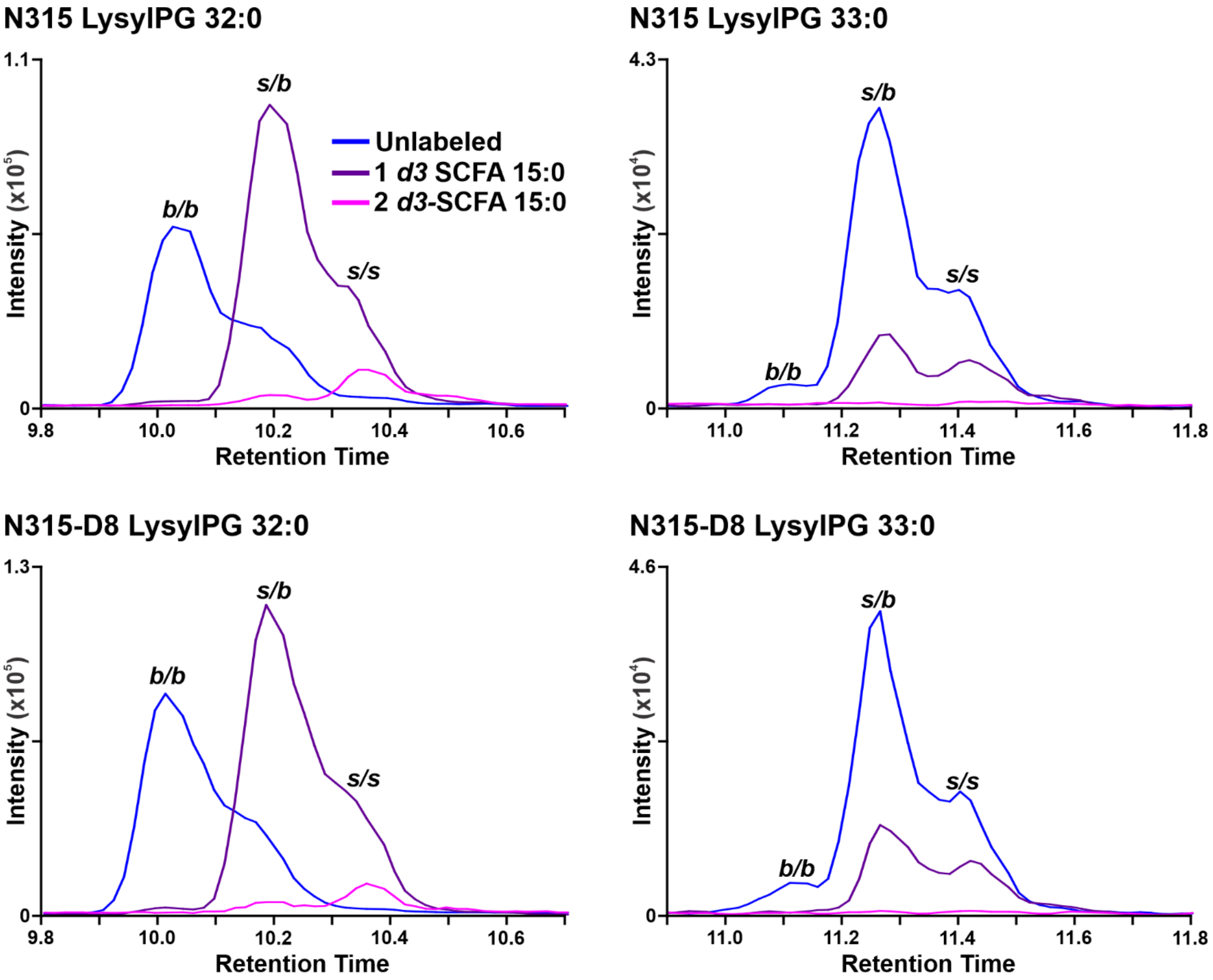


**Figure S12.** Extracted ion chromatograms of LysylPGs from N315 and N315-D8 grown in TSB supplemented with $d_3$-SCFA 15:0.

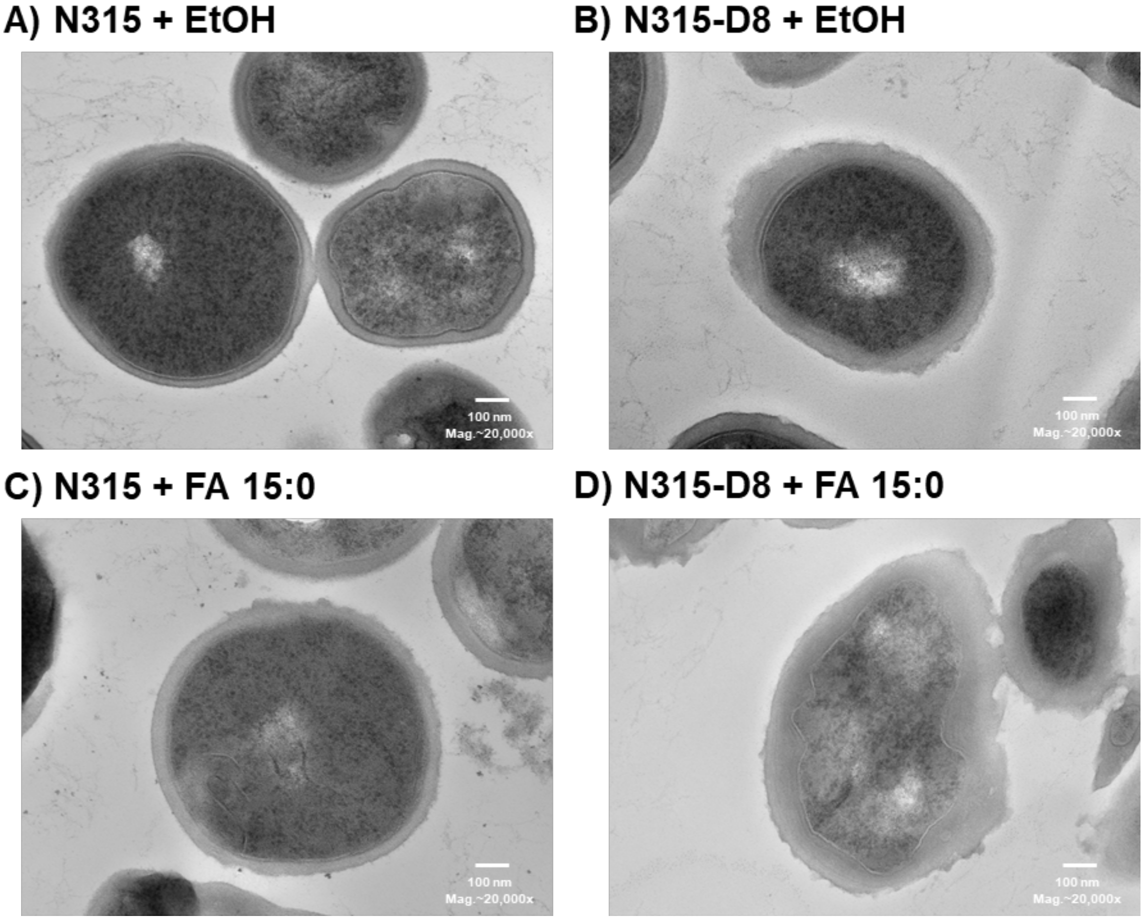


**Figure S13.** TEM images of N315 and N315-D8 cells that were treated with ethanol (as a vehicle control), or SCFA 15:0 in ethanol.

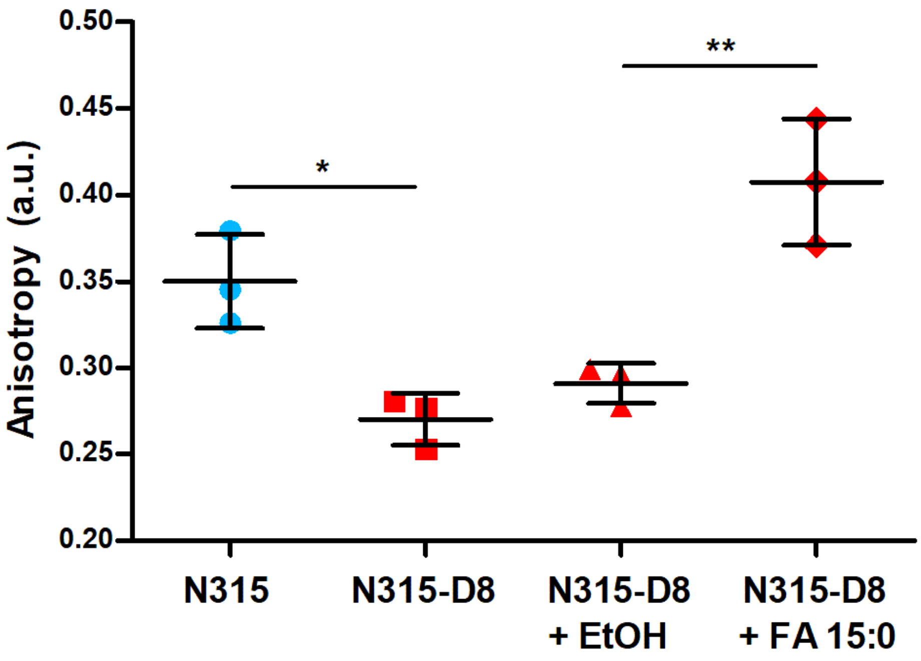


**Figure S14.** Anisotropic values of N315 and N315-D8 grown in TSB and N315-D8 supplemented with ethanol as a control and SCFA 15:0.

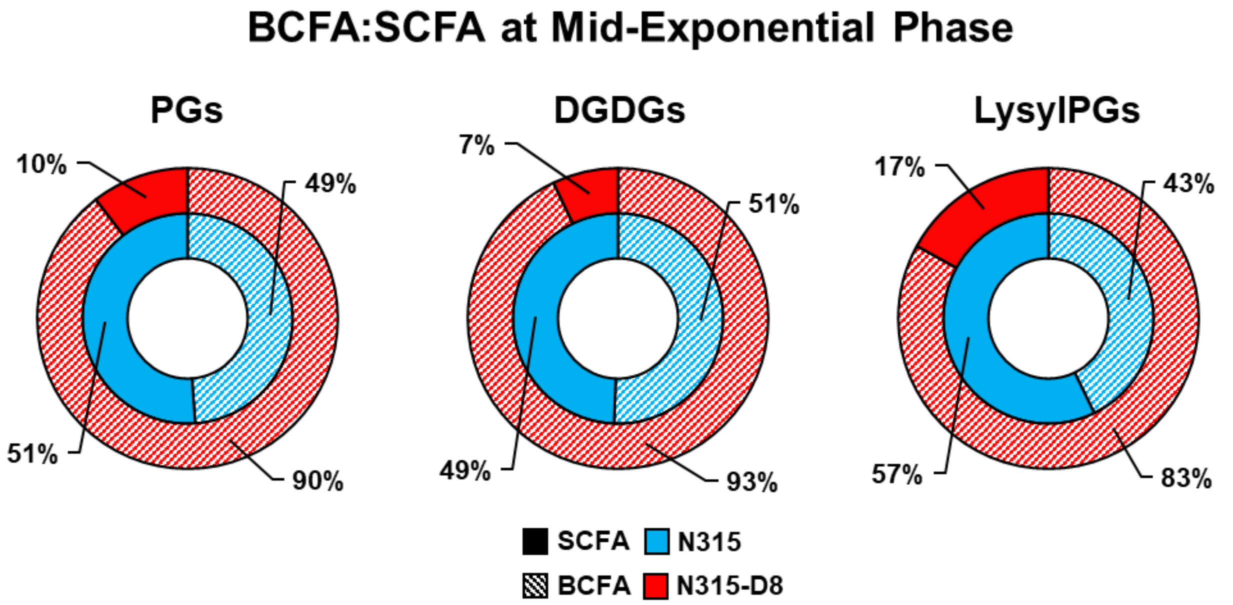


**Figure S15.** Distribution of BCFAs and SCFAs in PGs, DGDGs, and LysylPGs at mid-exponential phase in N315 and N315-D8.

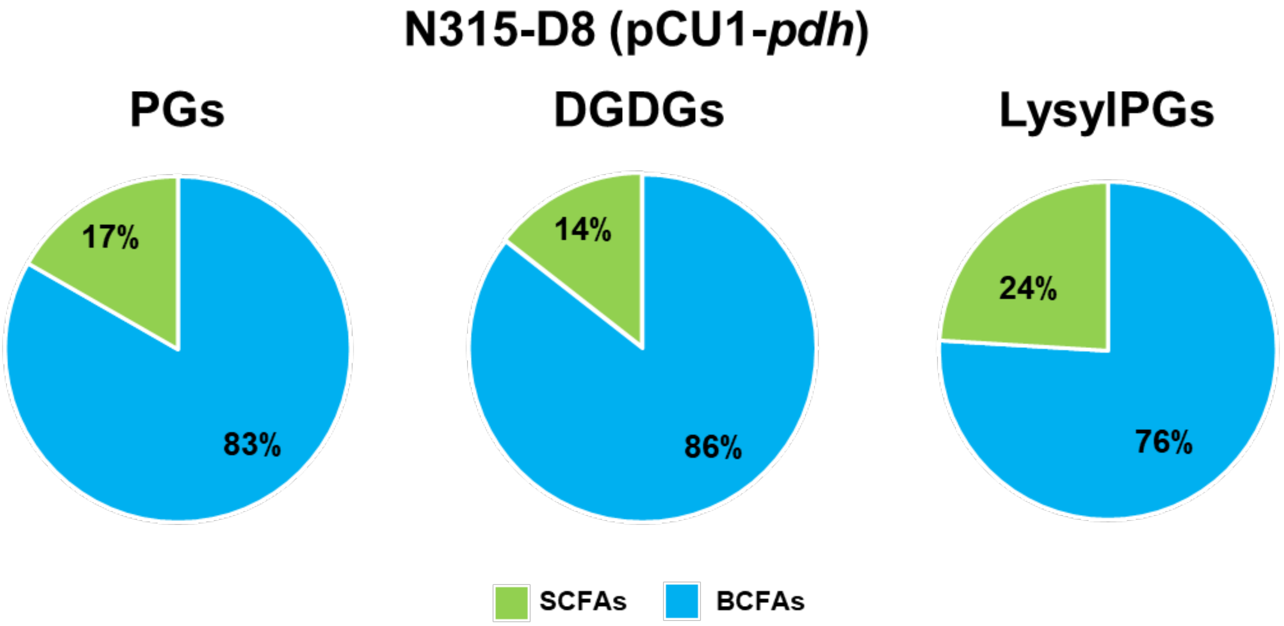


**Figure S16.** Pie charts of BCFA:SCFA for PGs, DGDGs, and LysylPGs in N315-D8 (pCU1-*pdh*).

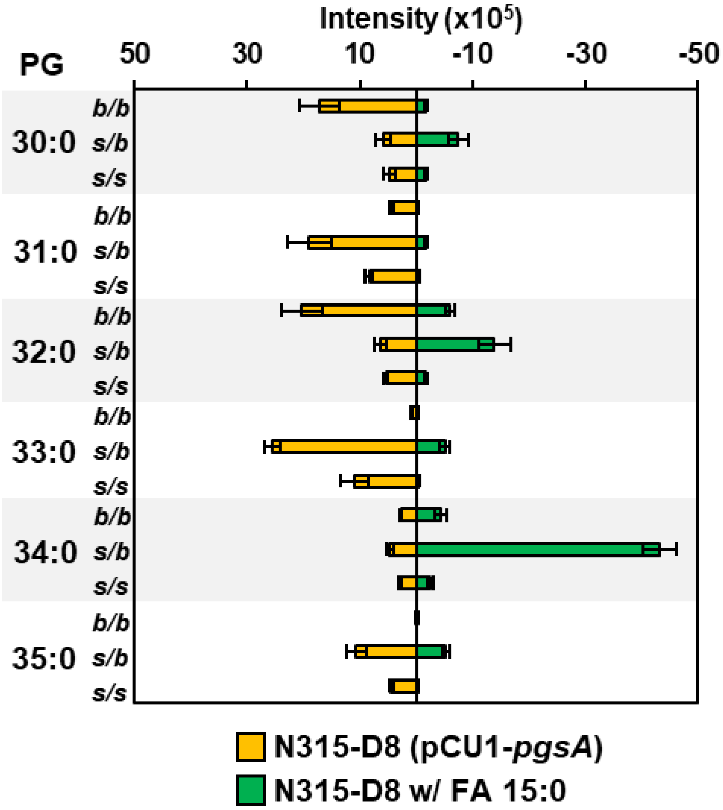


**Figure S17.** Distribution of BCFAs and SCFAs within PGs of N315-D8 (pCU1-*pgsA*) and N315-D8 grown in TSB supplemented with SCFA 15:0.

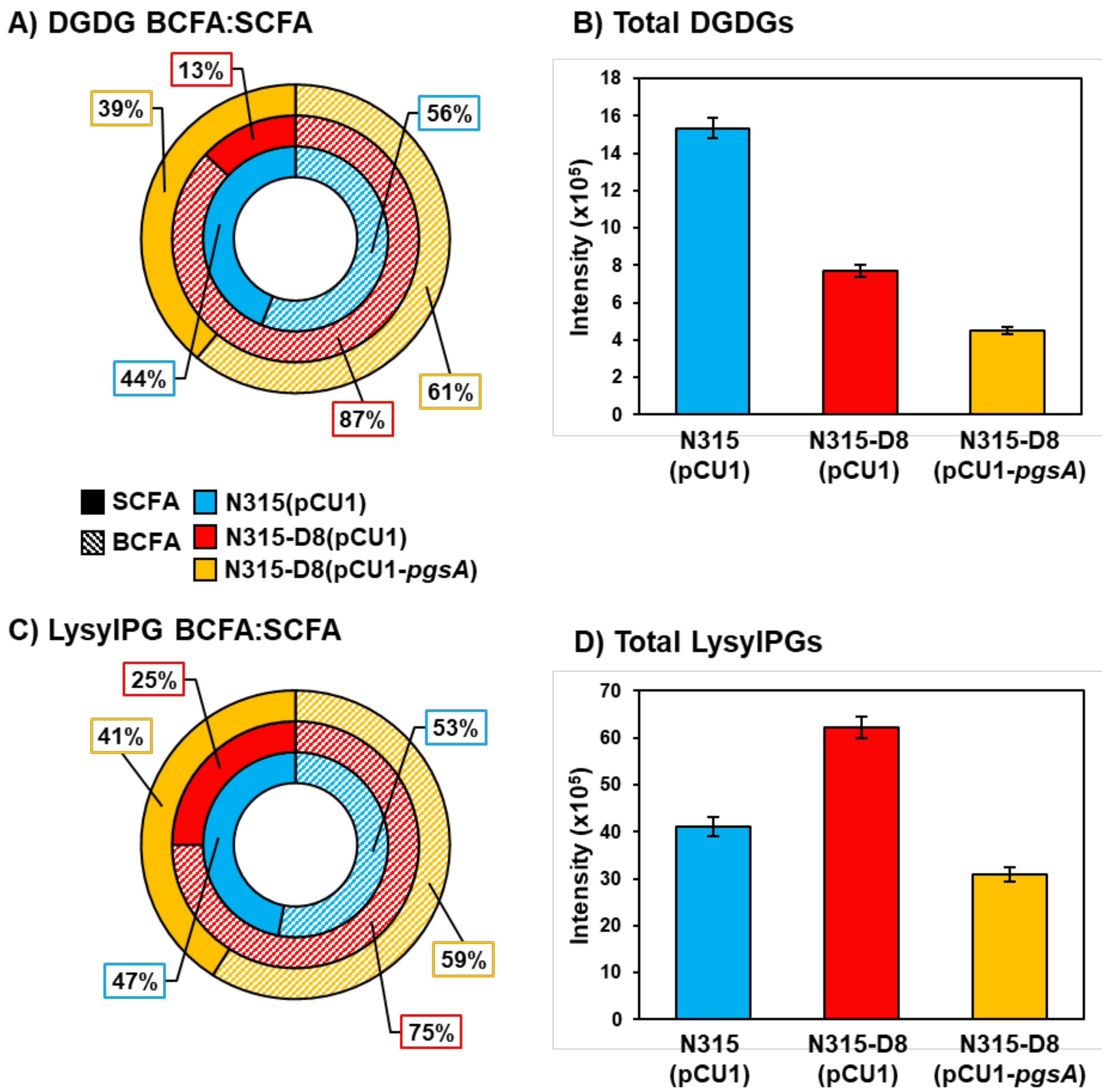


**Figure S18.** BCFA:SCFA ratios for A) DGDGs and C) LysylPGs, along with their total abundances (B,D), in N315 (pCU1), N315-D8 (pCU1) and N315-D8 (pCU1-*pgsA*).

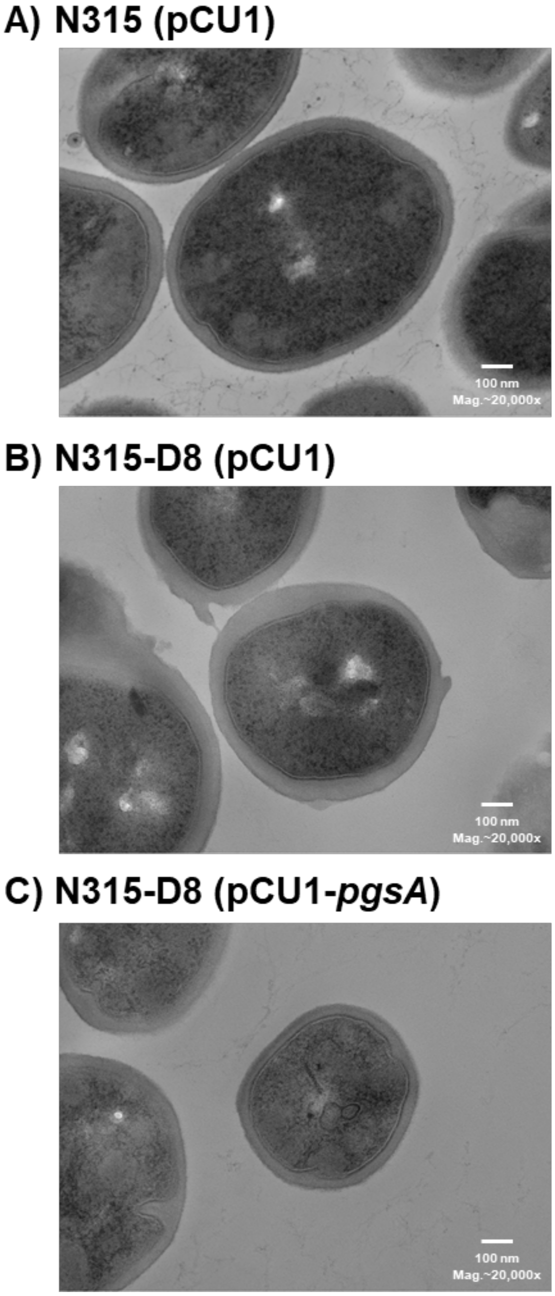


**Figure S19.** Representative TEM images of A) N315 (pCU1), B) N315-D8 (pCU1), and C) N315-D8 (pCU1-*pgsA*).

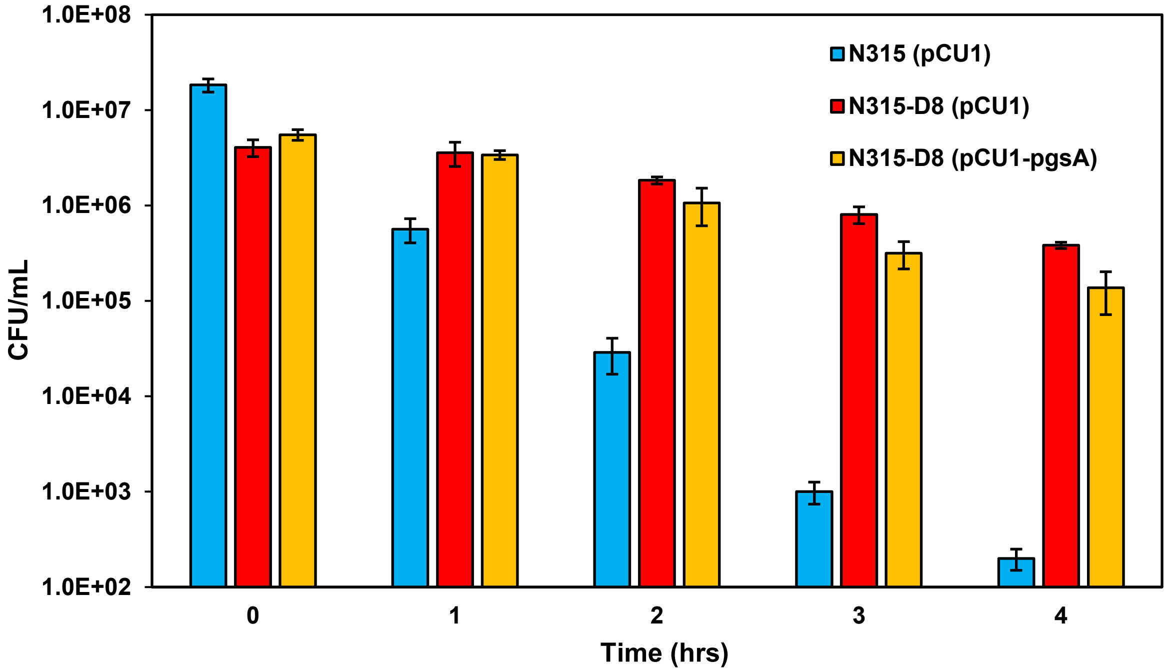


**Figure S20.** Number of cells (in CFU/mL) killed over 4 hours of daptomycin exposure (160 μg/mL in TSB) for N315 (pCU1), N315-D8 (pCU1) and N315-D8 (pCU1-*pgsA*).

**Table S2.** Daptomycin Minimum Inhibitory Concentrations.

| **Strain** | **Daptomycin MIC* (μg/mL)** |
|---|---|
| N315 | 0.5 |
| N315(pCU1) | 0.5 |
| N315-D8 | 8 |
| N315-D8(pCU1) | 8 |
| N315-D8(pCU1-*pdh*) | 8 |
| N315-D8(pCU1-*pgsA*) | 4 |

*Determined in Mueller-Hinton broth containing 30 mg/L $CaCl_2$.
